# Compression of functional gradients from rest to naturalistic processing: Moderation effect of age

**DOI:** 10.1101/2025.04.11.648387

**Authors:** Shuer Ye, Jasper Christian Mostert, Robin Pedersen, Xiaqing Lan, Lei Zhao, Alireza Salami, Maryam Ziaei

**Author notes:** Correspondence should be addressed to Maryam Ziaei, or Shuer Ye.

## Abstract

The functional cortical hierarchy of the human brain, a fundamental principle of brain organization, has been extensively characterized during resting state for healthy younger adults. However, functional re-organization during naturalistic settings, such as movie-watching, and its alterations across the adult lifespan remains poorly understood. Using resting-state and movie fMRI data from two large datasets, Cam-CAN (N=416) and DyNAMiC (N=156), this study conducted a comprehensive comparison of brain organization across two states. We identified a robust reorganization with compression of functional gradients from rest to movie-watching states, which is mediated by changes in functional integration and segregation of brain networks. The extent of compression from rest to movie was significantly greater in older adults and predicted worse cognitive performance among the elderly population. Our findings provide novel insights into how macroscale brain hierarchy is reorganized during naturalistic processing, and how this reorganization during aging impacts cognitive processes, offering a deeper understanding of the neural basis of aging and cognition.

## Introduction

Understanding how the brain integrates meaningful real-world stimuli to generate abstract cognition is pivotal for uncovering the neural foundations of human behavior (Chang et al., 2021; Y. Zhang et al., 2021). Current theories suggest that sensory inputs are integrated into higher-order cognitions along a cortical hierarchy spanning from unimodal to transmodal cortex (Bassett & Sporns, 2017; Bernhardt et al., 2022). A novel way to characterize the cortical hierarchy is through functional gradients, which map high-dimensional functional connectivity profiles onto low-dimensional manifolds, and provide a spatial framework for large-scale brain networks (Huntenburg et al., 2018; Jung et al., 2022; Knodt et al., 2023). A dominant sensorimotor-association (S-A) gradient has been consistently identified in resting state, spanning from sensorimotor regions to association regions(Margulies et al., 2016). This gradient has been robustly observed across populations and species (Tong et al., 2022; Valk et al., 2022). Other gradient including the visual-sensory (V-S) axis, which anchored in unimodal regions (Lei et al., 2023; Margulies et al., 2016), and the modulation-representation (M-R) axis, which spans from attention-related regions to sensory and association regions, have been also identified (Katsumi et al., 2022, 2023).

While the functional organization has been extensively characterized in resting-state studies, our understanding of how this hierarchy shifts in response to external stimuli remains limited. Recent works highlighted the reorganization of functional gradients across different contexts and cognitive states(Ito & Murray, 2023; Su et al., 2024). Reorganization occurs during acute pharmacological manipulation and in response to different tasks that reflect changes in the brain state, such as working memory load or sensorimotor adaptation (Gale et al., 2022; Girn et al., 2022; H. Zhang et al., 2022). The functional reorganization, thus, appears to reflect cognitive and attentional dynamics in macroscale brain organization during task engagements (Brown et al., 2022). However, controlled tasks often lack sufficient ecological validity (Finn, 2021). Naturalistic paradigms, such as movie-watching, embed rich, multimodal sensory information, driving continuous whole-brain activity akin to real-life experiences (Meer et al., 2020). A recent study revealed distinct gradients during movie-watching, likely reflecting dynamic sensory-cognitive interactions elicited by naturalistic stimuli (Samara et al., 2023). Yet, the reorganization from rest to movie-watching remains unexplored since functional gradients are typically estimated separately for each state.

The organization of functional gradients is closely linked to the balance between integration and segregation of functional networks, which reflect global communication and local specialization, respectively(Dong et al., 2024; Seguin et al., 2023). Developmental studies indicated that integration in sensory areas and segregation in association areas drive changes in functional gradients, supporting neurocognitive maturation (Pines et al., 2022; Y. Xia et al., 2022). Thus, striking a balance between specialized processing with global communication can ultimately support high-order cognitive functions (Senkowski & Engel, 2024; R. Wang et al., 2021). Naturalistic paradigms elicit enhanced integration between sensory and association areas(Ren et al., 2018; J. Wang et al., 2017) and improve sensitivity in predicting cognitive performance (Esmaeili et al., 2025; Finn & Bandettini, 2021). Therefore, gradient alterations during naturalistic stimuli likely reflect dynamic functional reorganization that balances between network integration and segregation.

Normal aging is associated with widespread changes in functional dynamics, with network dedifferentiation – marked by increased between-network and decreased within-network connectivity – leading to reduced functional specialization (Tsvetanov et al., 2016, 2018; Wu & Hoffman, 2024). This loss of functional specialization may underlie impairments in cognitive functions, such as working memory and cognitive flexibility (Nordin et al., 2024; Pedersen et al., 2023). Emerging evidence further links homogeneous connectivity patterns to cognitive decline, particularly in cortical regions with high tau deposition, a key pathological marker of Alzheimer’s disease (Ottoy et al., 2024). In older adults, a reduced hierarchical organization observed during resting state has also been associated with cognitive decline (Bethlehem et al., 2020). Despite these studies, it remains unclear how the functional organization varies across different states and whether these variations drive cognitive decline in aging. The dynamic reorganization across states reflects the brain’s adaptability in maintaining efficient information processing in response to task demands(Bola & Sabel, 2015; Vatansever et al., 2015). Therefore, unraveling how cortical gradients reorganize during state transitions may provide deeper insights into the mechanisms of brain aging, complementing single-state investigations and developing a more comprehensive framework for understanding the aging brain’s functional adaptability (Mooraj et al., 2025).

To bridge these gaps, this study aims, for the first time, to characterize alterations in cortical gradients during the transition from rest to movie-watching, investigate the mechanisms underlying functional reorganization, and examine how age influences this reorganization and its impact on cognitive performance. Two independent datasets - the Cambridge Centre for Ageing and Neuroscience (Cam-CAN) dataset and the DopamiNe, Age, connectoMe, and Cognition (DyNAMiC) dataset - were used to address our objectives (**Fig.1A**). We hypothesize that: (1) gradients compress during transitions from rest to movie-watching state; (2) functional integration and segregation drive this state-dependent gradient compression; (3) age exacerbates state-driven gradient compression along the S-A axis; and (4) gradient compression could predict cognitive performance, with this effect being most pronounced in older populations.

**Figure 1.**
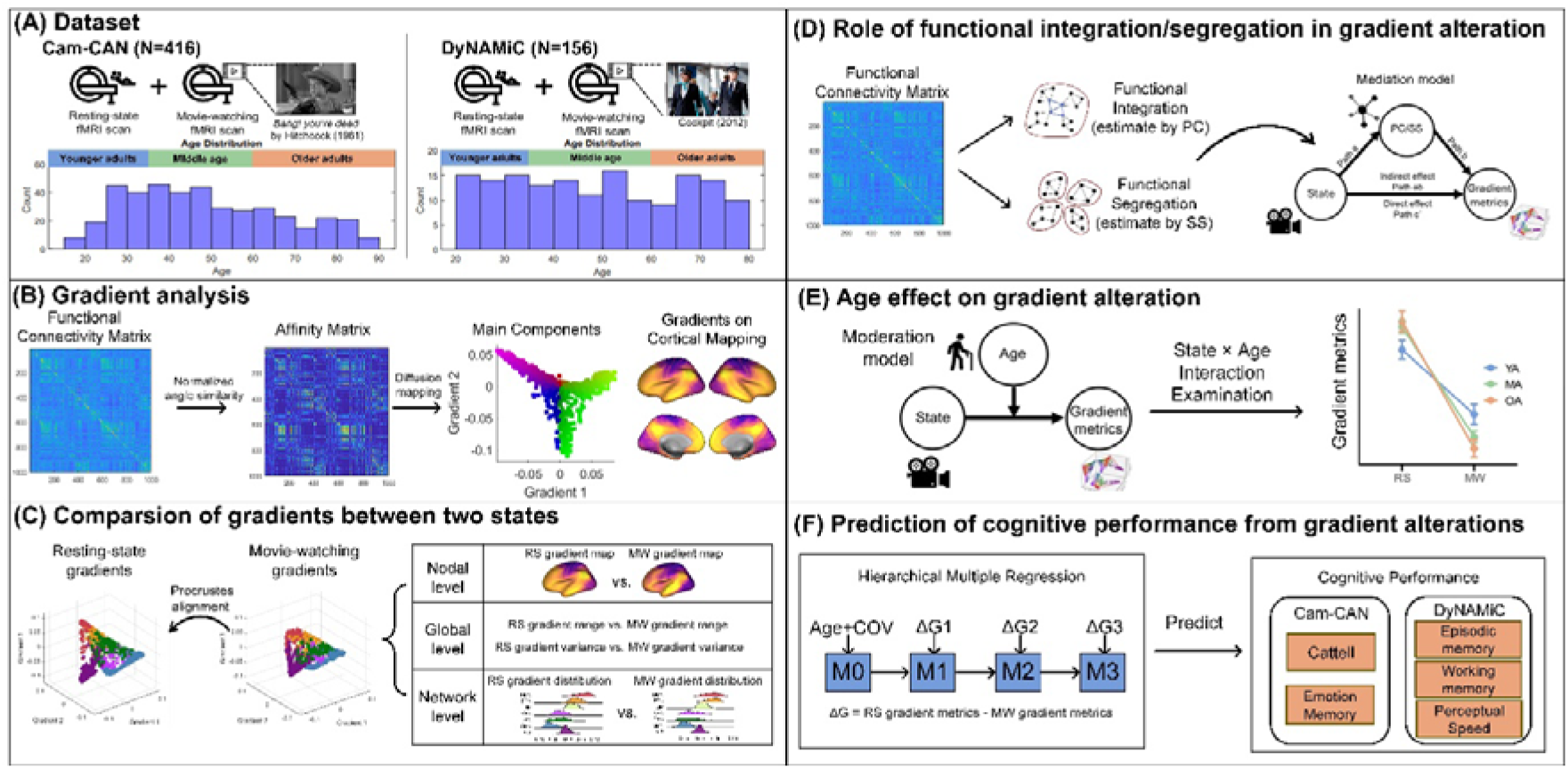
The workflow of the study. **(A)**Two large independent datasets spanning the lifespan including both resting-state and movie-watching fMRI were utilized. **(B)** Gradient analysis was applied to the r-z normalized and thresholded functional connectivity matrix. Diffusion mapping embedding was used to detect the gradient manifolds from the affinity matrix. Main gradients were identified based on explained variances. **(C)** The individual-level resting-state and movie-watching gradients were aligned to the resting-state group template to make the gradients of two states comparable. The gradients reorganization was comprehensively examined by comparing resting-state gradients and movie-watching gradients at nodal, global and network level. **(D)** Functional integration and segregation of the brain networks were estimated and compared between two states. Mediation models were constructed to test their role in the relationship between state and global gradient metrics. **(E)** Moderation models were constructed to examine the age effect on gradient alteration. **(F)** Hierarchical regression models were constructed to predict cognitive performance by adding gradient alteration parameters in a stepwise manner. RS=resting state; MW=movie watching; PC= participation coefficient; SS=system segregation; YA=younger adults’ group; MA=middle-age group; OA=older adults’ group. COV=covariance.

## Method

### Cam-CAN dataset

Two independent datasets were used to test our hypotheses. The first dataset taken from the publicly available Cam-CAN project Stage 2(Taylor et al., 2017). We included 649 participants who had completed both resting-state and movie-watching fMRI scans. After excluding participants with large head motion (average framewise displacement > 0.25 mm; Power et al., 2012) in either run, data from 416 participants (208 females, mean age: 48.77 ± 18.18 yrs, age range: 18.50-88.92 yrs) entered subsequent analyses. To index cognitive ability, measures of fluid intelligence and emotional memory performance were included in analyses. Fluid intelligence was assessed using the Cattell Culture Fair Intelligence Test, while emotional memory was measure with a computerized emotional memory task yielding three memory scores: priming, recognition, and recollection. Briefly, during training phase, participants viewed a background image with an emotional valence (neutral, negative or positive), followed by a foreground object and were instructed to imagine a story linking the two. In the subsequent testing phase, memory was assessed across three domains: (1) priming, by identifying a visually degraded object that was previously presented; (2) recognition, by rating confidence in whether the object had been presented earlier; and (3) recollection, by recalling the valence of the background image and verbally describing its content. Accuracy for each memory domains was quantified using the d ′ measure of discriminability, with higher d ′ scores indicate better performance. Further details of this task can be found in previous work (Shafto et al., 2014). Note that of the 416 participants in the original sample, 402 participants completed the fluid intelligence task, and 202 completed the emotional memory task; thus, analyses involving these measures were conducted on the respective subsamples. Demographic information of for the full sample of Cam-CAN is provided in **Tab.1**.

**Table 1.** Demographic information of Can-CAN dataset (N=416).

|  | Younger Group<br>[age<35yrs, N=113] | Mid-Age Group<br>[35yrs-60 yrs, N=185] | Older Group<br>[age> 60 yrs, N=118] | Overall<br>[N=416] |
| --- | --- | --- | --- | --- |
| Age (yrs) | 28.07 ± 4.50 | 28.07 ± 4.50 | 72.79 ± 8.30 | 48.77 ± 18.18 |
| Sex (N) |  |  |  |  |
| Female | 63 (56%) | 92 (50%) | 53 (49%) | 208 (50%) |
| Male | 50 (44%) | 93 (40%) | 65 (51%) | 208 (50%) |
| BMI | 23.15 ± 3.16 | 24.73 ± 3.79 | 25.49 ± 3.09 | 24.51 ± 3.55 |
| Head motion (mm) |  |  |  |  |
| Resting-state | 0.14 ± 0.05 | 0.16.8 ± 0.06 | 0.21 ± 0.05 | 0.17 ± 0.06 |
| Movie-watching | 0.12 ± 0.05 | 0.15 ± 0.06 | 0.16 ± 0.06 | 0.15 ± 0.06 |
| TIV (mm<sup>3</sup>) | 1498.46 ± 162.01 | 1521.06 ± 157.53 | 1491.96 ± 151.92 | 1506.31 ± 157.10 |
| Income |  |  |  |  |
| <£18000 | 16 (14.2%) | 12 (6.5%) | 30 (25.4%) | 58 (13.9%) |
| £18000-30999 | 28 (24.8%) | 27 (14.6%) | 40 (33.9%) | 95 (22.8%) |
| £31000-51999 | 34 (30.1%) | 53 (28.7%) | 27 (22.9%) | 114 (27.4%) |
| £52000-100000 | 25 (22.1%) | 69 (37.3%) | 17 (14.4%) | 111 (26.7%) |
| >£100000 | 10 (8.9%) | 24 (13.0%) | 4 (3.3%) | 38 (9.1%) |
| Education |  |  |  |  |
| Degree | 82 (72.6%) | 142 (76.8%) | 55 (46.6%) | 279 (67.1%) |
| A-level | 17 (15.0%) | 27 (14.6%) | 28(23.7%) | 72 (17.3%) |
| O-level | 14 (12.4%) | 15 (8.1%) | 21(17.8%) | 50 (12.0%) |
| None | 0 (0%) | 1 (0.5%) | 14 (11.9%) | 15 (3.6%) |

Note: BMI= body mass index; TIV= total intracranial volume.

The Cam-CAN MRI data was collected by a 3T Siemens TIM Trio scanner with a 32-channel head coil. The resting-state fMRI data were acquired using an echo planar imaging (EPI) sequence: voxel size = 3×3×4.44 mm^3^; 32 axial slices, TR = 1970 ms; TE = 30 ms; FA = 78 °; FOV: 192×192 mm^2^. The resting-state was in eyes-closed condition and lasted for 520 seconds. The movie-watching functional data were acquired using a multi-echo EPI sequence: voxel size = 3×3×4.44 mm^3^; 32 axial slices; TR = 2470 ms; TE = 9.4, 21.2, 33, 45, 57 ms; FA = 78°; FOV: 192×192 mm^2^. During the movie, participants watched a 480-second clip from the “*Bang! You’re Dead*” movie. The Cam-CAN imaging data were preprocessed using the Automatic Analysis (AA) pipeline provided by the Cam-CAN project. Full details of the pipeline are described in the report by Taylor and colleagues(Taylor et al., 2017). Briefly, functional images were corrected for motion and slice timing, co-registered to the T1-weighted structural image, and normalized into standard MNI space at 3mm resolution. On top of the AA-preprocessed data, additional preprocessing was performed. This included linear detrending, nuisance regression (white matter signal, cerebrospinal fluid signal, and 12 head motion parameters), temporal filtering with a band-pass of 0.009-0.08 Hz, and spatial smoothing with a 4 mm full-width-at-half-maximum (FWHM) Gaussian kernel using Nilearn library in Python. The first 10 volumes were discarded to ensure magnetization equilibrium.

### DyNAMiC dataset

The second dataset is a part of the DopamiNe, Age, connectoMe, and Cognition (DyNAMiC) project. Full details of the project can be found in Nordin et al (Johansson et al., 2023; Nordin et al., 2022). For the first wave of DyNAMiC, 180 healthy participants with age ranges between 20-80 years old were recruited. All the participants underwent a 12-minute eye-open resting-state scan and a 12-minute naturalistic movie-watching scan. After excluding participants with large head motion (average framewise displacement > 0.25 mm) in either the rest or movie run, data from 156 participants (82 females, mean age: 48.03 ± 17.25 yrs, age range: 20-77 yrs) entered the subsequent analyses. Demographic information of for the full sample of DyNAMiC is provided in **Tab.2**. A battery of cognitive tests was used to assess three key cognitive domains: episodic memory, working memory, and perceptual speed, all of which were included in our analyses.

**Table 2.** Demographic information of DyNAMiC dataset (N=156).

|  | Younger Group<br>[age<35 yrs, N=44] | Mid-Age Group<br>[35yrs-60 yrs, N=66] | Older Group<br>[age>60 yrs, N=46] | Overall<br>[N=156] |
| --- | --- | --- | --- | --- |
| Age (yrs) | 26.95 ± 4.34 | 47.11 ± 7.28 | 69.50 ± 4.91 | 48.03 ± 17.25 |
| Sex (N) |  |  |  |  |
| Female | 22 (50%) | 35 (53%) | 25 (54%) | 82 (53%) |
| Male | 22 (50%) | 31 (47%) | 21 (46%) | 74 (47%) |
| BMI | 24.60 ± 4.18 | 25.60 ± 3.66 | 25.36 ± 4.56 | 25.27 ± 4.11 |
| Head motion (mm) |  |  |  |  |
| Resting-state | 0.11 ± 0.05 | 0.13 ± 0.05 | 0.17 ± 0.07 | 0.14 ± 0.06 |
| Movie-watching | 0.05 ± 0.04 | 0.13 ± 0.06 | 0.15 ± 0.06 | 0.13 ± 0.06 |
| TIV (mm<sup>3</sup>) | 1627.60 ± 183.18 | 1567.71 ± 159.45 | 1531.25 ± 167.00 | 1573.81 ± 157.10 |
| Education (yrs) | 15.03 ± 2.07 | 15.86 ± 3.37 | 14.48 ± 3.99 | 15.22 ± 3.31 |

Note: BMI= body mass index; TIV= total intracranial volume.

The imaging data were collected with a 3T Discovery MR 750 scanner (General Electric), equipped with a 32-channel phased-array head coil. Functional images were sampled using a T2*-weighted single-shot EPI sequence: voxel size = 1.95×1.95×3.4 mm^3^; 37 axial slices; TR = 2000 ms; TE = 30 ms, FA = 80°, and FOV = 250 × 250 mm. Ten dummy scans were collected at the start of the sequence. The acquisition for both functional runs lasted 12 minutes, resulting in a total of 360 volumes. During the resting-state scan, the participants were instructed to keep their eyes open and focus on a white fixation cross presented on a black background. During the movie-watching scan, participants viewed and listened to a 12-minute video consisting of selected and chronologically ordered sections from the Swedish movie “Cockpit”.

A similar preprocessing protocol of the functional images was applied in DyNAMiC as in Cam-CAN. These steps included removal of dummy scans, co-registration to the T1-weighted structural image and normalized into standard MNI space at 2 mm resolution, linear detrending, nuisance regression (white matter signal, cerebrospinal fluid signal, and 12 head motion parameters), temporal filtering with a band-pass of 0.009-0.08 Hz, and spatial smoothing with a 4 mm FWHM Gaussian kernel.

### Gradient analysis

The Schaefer atlas with 1000 parcels was used to compute ROI-wise time series for each subject (Samara et al., 2023; Schaefer et al., 2018). These 1000 regions can be assigned to seven spatially independent functional networks: visual network (VIS), somatomotor network (SMN), dorsal attention network (DAN), ventral attention network (VAN), limbic network (LIM), frontoparietal network (FPN), and default mode network (DMN). Functional connectivity was calculated by computing the Pearson correlation between the time courses of each pair of parcels followed by Fisher’s z-transformation to create subject-specific 1000 × 1000 functional connectivity matrices for the resting and movie-watching states, respectively.

To estimate the functional gradient, the connectivity matrices were thresholded row-wise with the top 10% of the connections retained. Normalized angle similarity was applied to generate symmetric affinity matrices without negative values to denote the similarity of connectivity profiles between brain regions(Chakraborty et al., 2024; Huang et al., 2023). Diffusion map embedding, a nonlinear dimensionality reduction algorithm, was used to compute the gradient components explaining the most variance in functional connectivity profiles(Coifman et al., 2005). For the diffusion embedding, the manifold learning parameter α, which controls the influence of sampling point density on the manifold, was set to the default value of 0.5 in accordance with previous gradient studies(Margulies et al., 2016; Samara et al., 2023). To ensure comparability between individuals as well as states, we used Procrustes rotation to align individual gradient maps from both the resting and movie-watching states to a group-level gradient template generated from the averaged resting-state functional connectivity matrix (Langs et al., 2015). For the sake of completeness, although it’s not the focus of this study, we estimated the independent movie-watching gradients (i.e., individual movie-watching gradients were Procrustes aligned to the group-average movie-watching template instead of the resting-state template). These independent movie-watching gradients were presented in **SFig.1**. These procedures (**Fig.1B**) were implemented using the BrainSpace toolbox (Vos de Wael et al., 2020) in MATLAB2023b.

The functional gradients are ranked based on the amount of variance explained in descending order. The first three gradients were selected for further analysis as they explained most of the variance (for resting state, the first three gradients explained 38.44% and 38.05% of total variance in the Cam-CAN and DyNAMiC, respectively). The gradient score for each region represents its relative position in the functional hierarchy. Two global gradient metrics were computed for each gradient: range (i.e., the absolute differences between maximum and minimum gradient scores) and variance (i.e., the standard deviation of the gradient scores). Specifically, gradient range represents the degree of dissimilarity in connectivity patterns between regions situated at the opposite poles of the gradient manifold, while gradient variance reflects the heterogeneity in the connectivity profile across regions.

To better characterize the topography of the gradients, we conducted a meta-analysis to identify the functional and anatomical terms associated with regions of the gradient poles (i.e., top 10% and bottom 10% of the gradient scores) of the selected gradients by using the NeuroSynth database. Specifically, the upper and lower 10% gradient maps were used as input to the analysis, and the output are feature terms with *r*-values. The *r*-value represents the strength and direction of the association between the spatial pattern the gradient map and the meta-analytic activation maps from the NeuroSynth database (Yarkoni et al., 2011). The terms were then ordered in a descending order based on the *r*-value and visualized in a word cloud to indicate the relevance of these terms.

### Functional integration and segregation

The extent of functional integration was assessed using a measure of the participation coefficient (PC), a graph theoretical metric that measures how one brain regions connect to other brain regions of different networks (Power et al., 2013). The calculation of PC was performed using unthresholded Fisher r-z transformed connectivity matrices. These matrices were subsequently binarized to create undirected and unweighted graphs, considering only positive connectivity values and applying a sparsity threshold of 0.1 (i.e., only the 10% most stronger connections were retained). Based on the pre-defined seven networks of the Schaefer atlas, the participation coefficient for each region (node) can be calculated as the following formula:

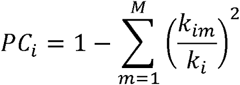

Based on the binary graph, *m* represents a specific module within the set of modules *M*. The term *k_i_* denotes the total number of connections associated with node *i* across the entire graph, while *k_im_* refers to the number of connections between node *i* and module *m*. For each participant, the nodal PC were averaged across brain regions to quantify the global functional integration, and higher global PC indicates increased function integration.

The degree of functional segregation was evaluated using a measure of system segregation (SS) (Chan et al., 2014). SS quantifies the relative strength of within-network connectivity in relation to between-network connectivity, as expressed by the following formula:

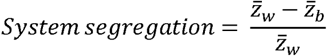

Based on un-thresholded Fisher-transformed connectivity matrices that take only positive connections, the within-network connectivity strength, *Z_w_*, was calculated as the average of connectivity weights between nodes within the same network, and between-network connectivity strength, *Z_b_*, was calculated as mean of connectivity weights between nodes of a given network and nodes in other networks. Here, higher SS value indicates increased functional segregation of the brain networks.

### Statistical Analyses

During statistical analyses, we considered sex, education level, income, body mass index (BMI), total intracranial volume (TIV), and head motion as covariables of no interest, thus they were regressed out. The TIV of each subject was estimated after the brain extraction from the T1 image using the Computational Anatomy Toolbox (CAT12) (Gaser et al., 2024). It should be noted that we only included sex, BMI, TIV, education level, and head motion as a confound in statistical analyses of the DyNAMiC dataset as income information was not available in this dataset. Benjamini-Hochberg False Discovery Rate procedure (FDR correction) was applied in multiple comparisons to control for the risk of false positives.

### Assessment of state-dependent difference in functional gradient

The state-dependent differences in each functional gradient were examined at regional, global and network levels, through performing a series of paired sample *t*-tests **(Fig.1C)**. First, we compared regional gradient scores across brain regions of each gradient between resting and movie-watching state. The resulting whole-brain t-map obtained from the difference analysis was then correlated with the resting-state gradient map to investigate the direction of gradient alterations at the regional level. To account for spatial autocorrelation, and to assess the significance of the spatial correlation between resting-state gradient map and the t-map of gradient difference between the two states, spin permutation tests were performed using the BrainSMASH toolbox (Burt et al., 2020) based on Python. We generated 10,000 surrogate brain maps with preserved spatial autocorrelation from the empirical t-map to produce a null distribution. The significance *p*_spin_ was obtained by comparing the observed *t-* maps with the surrogate maps. Subsequently, we compared the global gradient metrics (i.e., gradient range and variance) for each gradient to indicate the gradient alterations at the global level. Additionally, network gradient scores were calculated by averaging the gradient scores of all nodes within each network, and these were compared between the resting and movie-watching state to examine how network positions along the gradient axis shift from rest to movie-watching.

### Linking state-driven gradient alteration with functional integration and segregation

Paired-sample *t*-tests were performed to examine the state differences in network integration and segregation. We also assessed the association of functional integration and segregation with gradient metrics for each state using partial correlation analysis, controlling for the effect of age. Next, mediation analyses were conducted to determine whether state-driven gradient alterations were mediated by functional integration or segregation **(Fig.1D)**. Specifically, mediation models were constructed with state (i.e., a binary variable indicating either resting-state or movie-watching) as the independent variable, global PC or SS as the mediation variable (depending on whether it shows a significant difference between the two states), and gradient metrics as the dependent variable. Thus, the mediation model included the following paths: 1) Path a: the effect of the independent variable (i.e., states) on the mediator (i.e., functional integration/segregation); 2) Path b, the effect of the mediator on the dependent variable (i.e., gradient metrics), while controlling for the independent variable; 3) Path ab: The indirect effect of the independent variable on the dependent variable through the mediator, calculated as the product of Path a and Path b; and 4) Path c’: The direct effect of the independent variable on the dependent variable after accounting for the mediator. A significant indirect effect would indicate that the relationship between state and gradient metrics is partially or fully mediated by functional integration or segregation. The mediation analyses were conducted by using “lavaan” package in R version 4.4.1, controlling for the same covariates as mentioned above during the analyses.

### Moderation effect of Age on state-driven gradient alteration

To address our third hypothesis, linear regression models with state-by-age interactions were constructed to assess whether age moderates state-driven gradient difference using the following formula **(Fig.1E)**:

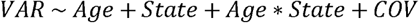

Here, VAR represents the gradient metrics (i.e., gradient range or variance) and COV represents covariables as same as mentioned above. This procedure was conducted by using the *lm* function in R version 4.4.1.

### Predict cognitive performance based on gradient compression

To investigate the association between cognitive ability and the degree of gradient compression, we first calculated the difference in gradient range and variance between resting-state and movie-watching by subtracting the movie-watching gradient metrics from the resting-state gradient metrics. Higher values of these six difference values (i.e., ΔG1-range, ΔG1-variance, ΔG2-range, ΔG2-variance, ΔG3-range, ΔG3-variance) indicate greater gradient compression. Next, hierarchical multiple regression (HMR) was applied to examine whether the extent of gradient compression could predict cognitive performance **(Fig.1F)**. This method allows estimation of the unique contribution of each set of predictors (i.e., compression for each gradient) in relation to the variance of the dependent variable while controlling for the effects of other covariates. For the first step, age and covariates were entered in the first step to construct a base model (M0), with cognitive performance as the dependent variable. Subsequently, the difference parameters of the three gradients were added as predictor variables into the model step by step, resulting in three models (i.e., Model1 [M1] with differences of G1 range and variance entered, Model2 [M2] with differences of G2range and variance entered, and Model3 [M3] with differences of G3 range and variance entered) to predict cognitive performance. We further repeated the hierarchical multiple regression in three age groups (i.e., younger adults, middle-age, and older adults) to investigate the age-related changes in the relationship between gradient compression and cognitive performance.

## Results

### Compressed gradient pattern from rest to movie

The cortical topographic mapping and spatial distribution of the resting-state and movie-watching gradients from Cam-CAN dataset are presented in **Fig.2A**. The first three gradients (i.e., G1, G2, and G3) explained 38.44% of variance for resting-state (G1:14.32%; G2:12.74%; G3:11.37%), and 34.33% of variance for movie-watching (G1:14.27%; G2:10.91%; G3: 9.15%). The principal gradient (G1) is the visual-sensorimotor gradient (i.e., V-S axis), which reveals a hierarchy of unimodal systems spanning from visual to sensorimotor regions. The second gradient (G2) is the unimodal-transmodal gradient (i.e., S-A axis) with a hierarchical axis from visual and sensorimotor regions to the DMN areas. The third gradient (G3) primarily separates sensorimotor, insular, and temporal regions from the DMN, including the posterior cingulate cortex, temporoparietal junction, and medial prefrontal cortex, resulting in an insular-DMN axis. Results from the meta-analytic decoding analysis suggest functional and anatomical terms for each pole of gradients, demonstrating the characterization of the selected gradients (**SFig.2**).

**Figure 2.**
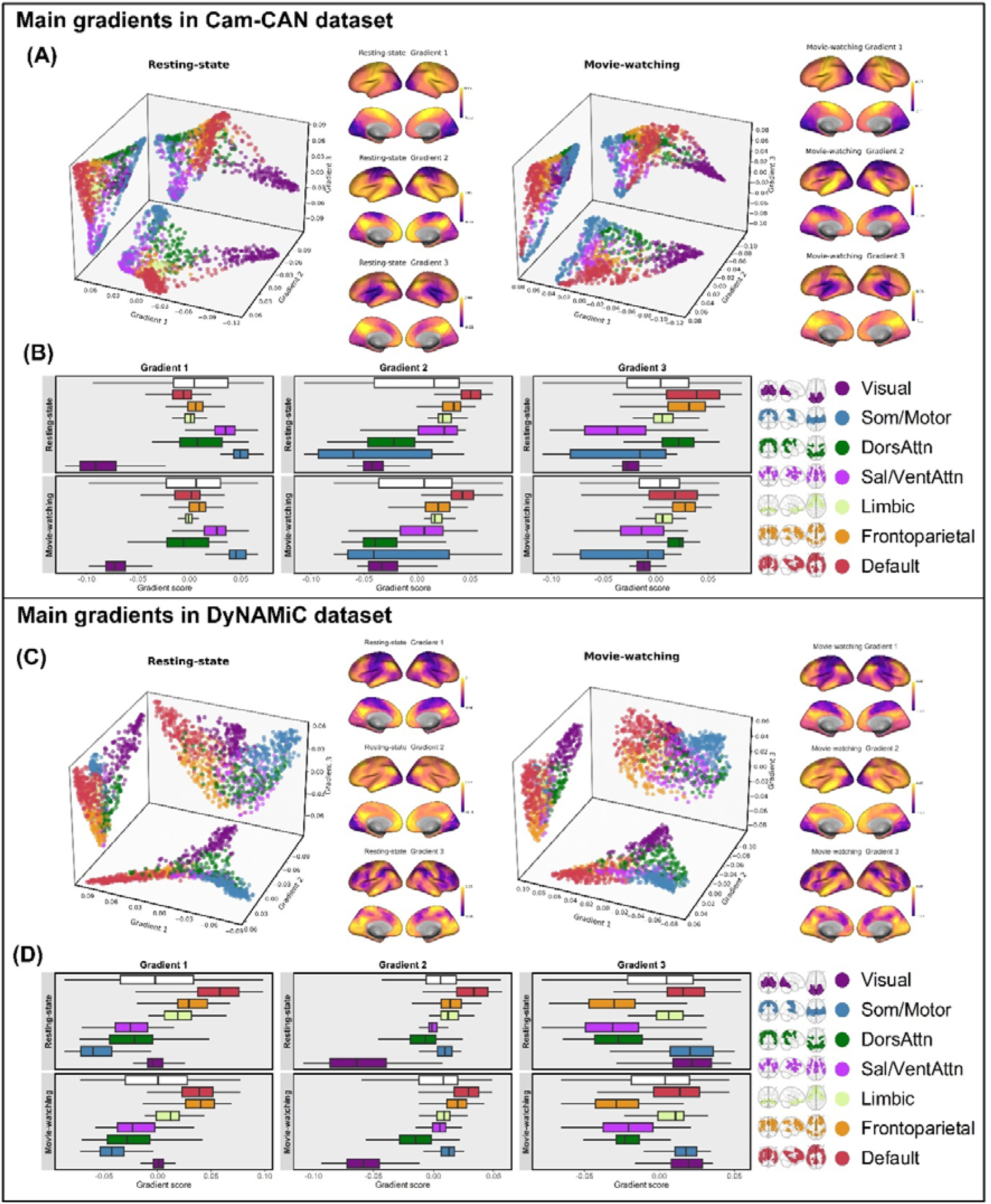
The first three functional gradients of resting and movie-watching states across the two datasets. **(A/C)** The 2D projection of the gradients in 3D space, gradient projections onto the cortical surface. **(B/D)** Boxplots of the gradient distribution. Scatterplots and boxplots of the gradients in resting and movie-watching states color-coded by parcel assignments of Yeo’s 7 functional networks. The white box indicates the overall gradient scores. The lower edge of the box represents the 1st quartile, the upper edge represents the 3rd quartile, and the line inside the box indicates the median of the gradient score.

To investigate our first hypothesis, whether gradients compress during transitions from rest to movie-watching state, we started with comparing gradient scores (i.e., eigenvector values denoting relative position of regions in the embedded space) across resting-state and movie-watching at the regional level and mapped the *t*-value to the cortex. We observed that areas with larger differences in gradient scores significantly overlapped with the two poles of the gradient axis (**Fig.3A**). Importantly, the positive pole (e.g., sensorimotor areas in G1, DMN subregions in G2 and G3) tends to decrease during the transition from rest to a movie-watching state, whereas the negative pole (e.g., visual areas in G1, sensorimotor areas in G2, and the insula in G3) tends to increase. This shift leads to a reduction in the range between the two gradient poles. Further correlation analysis revealed a significant positive spatial correlation between the *t*-values and the resting-state gradient score (G1: *r* = 0.527, *p_spin_* < 0.001; G2: *r* = 0.551, *p_spin_* < 0.001; G3: *r* = 0.708, *p_spin_* < 0.001), which confirms the trend of compression. These findings suggest that the gradient anchors converge toward the center when transitioning from rest to movie, indicating a pattern of state-driven gradient compression, in which the overall constellation of cortical hierarchy is reduced during movie watching.

**Figure 3.**
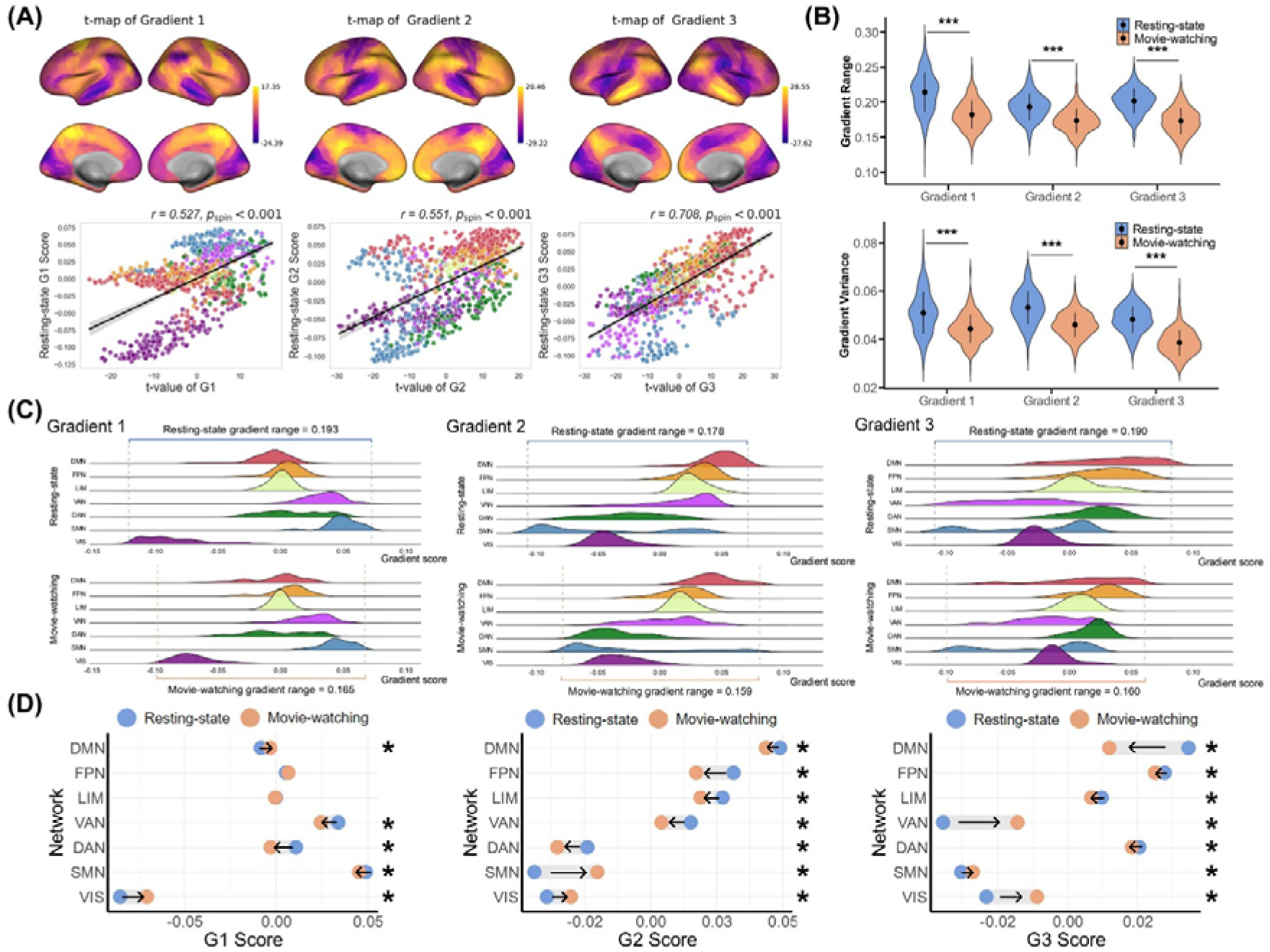
Regional, global, and network level gradient differences between resting state and movie watching: results from Cam-CAN dataset. **(A)** The t-map of the difference in the regional gradient score between two states, and the correlation between the t-map and resting-state gradient score. The significance of the correlation was estimated by using the spin permutation test. The shaded zone represents 95% confidence interval. **(B)** The violin plot illustrates that the gradient ranges and variances of the three gradients are higher during rest compared to naturalistic processing. **(C)** The distribution of the gradient score for Yeo’s 7 networks at two states. **(D)** The difference in network gradient score between two states. The dot represents the network gradient score, and the arrow indicates the moving direction of the network position on the gradient axis when transitioning from rest to movie watching. VIS=visual network; SMN=somatomotor network; DAN=dorsal attention network; VAN=ventral attention network; LIM=limbic network; FPN=frontoparietal network; DMN= default mode network. *: p<0.05; ***:<0.001.

Next, we compared the global gradient metrics of range and variance. The results of the paired *t*-test indicated an overall larger gradient range and variances of resting-state gradients compared with movie-watching gradients (G1 range: *t* = 18.596, *d* = 0.911, *p* < 0.001; G2 range: *t* = 16.788, *d* = 0.823, *p* < 0.001; G3 range: *t* = 23.156, *d* = 1.135, *p* < 0.001; G1 variance: *t* = 12.462, *d* = 0.611, *p* < 0.001; G2 variance: *t* = 17.409, *d* = 0.853, *p* < 0.001; G3 variance: *t* = 26.001, *d* = 1.274, *p* < 0.001, **Fig.3B**). The results revealed a global compression of the gradient when transitioning from the resting state to the movie-watching state, as further illustrated in **Fig.3C**. Moreover, comparisons analysis between the network-average gradient scores indicated significant changes between two states (**STab.1**). It was repeatedly observed in all three gradients that, when transitioning from rest to movie-watching, the network positions shift toward the middle of the gradient (**Fig.3D**). Taking G1 as an example, compared to the resting state, the average gradient scores of the VIS and DMN increased, while those of the DAN, VAN, and SMN decreased, converging toward the global average during movie-watching. To visualize the state-driven gradient compression, we displayed an animation of a 3D scatter plot (can be viewed in https://osf.io/p6cza) to show how resting-state gradients transform into movie-watching gradients.

For DyNAMiC dateset, the first three gradient explained 38.05% of variance for resting-state (G1:16.73%; G2:12.06%; G3:9.26%), and 36.04% of variance for movie-watching (G1:15.33%; G2:11.54%; G3: 9.17%). G1 represents a S-A axis, with its unimodal apex primarily located in sensorimotor areas and visual areas posited in the middle of the manifold. G2 also follows a unimodal-to-transmodal gradient along a hierarchical axis extending from visual regions to the association areas, forming a visual-nonvisual gradient. G3 aligns with the M-R axis, primarily distinguishing attention-related areas from visual, sensorimotor, and association regions **(Fig.2B)**. Among these main gradients, Cam-CAN G1 and DyNAMiC G2, Cam-CAN G2 and DyNAMiC G1 show the highest similarity (**SFig.3**).

The comparison between resting-state and movie-watching gradients in DyNAMiC similarly reveals significant gradient compression from resting state to movie-watching, consistent our observations in Cam-CAN dataset (**Fig.4**). Specifically, the positive pole (e.g., association areas in G1 and G2, unimodal areas in G3) exhibits a decrease, while the negative pole (e.g., sensorimotor areas in G1, visual areas in G2, and attention-related areas in G3) shows an increase during movie watching compared to rest. Moreover, the gradient state-difference t-maps are highly correlated with the resting-state gradient maps (G1: *r* = 0.694, *p_spin_* < 0.001; G2: *r* = 0.357, *p_spin_* < 0.001; G3: *r* = 0.626, *p_spin_* < 0.001). These findings indicate that the two poles of the main gradients move closer to the center when transitioning from rest to movie watching, suggesting a more compressed functional hierarchy during naturalistic processing, independent of dataset. The state-driven gradient compression was further supported by state-to-state comparisons at both the global and network levels. Resting-state global gradient range and variance are significantly greater than those of movie-watching state (G1 range: *t* = 12.159, *d* = 0.973, *p* < 0.001; G2 range: *t* = 11.198, *d* = 0.897, *p* < 0.001; G3 range: *t* = 12.035, *d* = 0.963, *p* < 0.001; G1 variance: *t* = 11.057, *d* = 0.885, *p* < 0.001; G2 variance: *t* = 6.538, *d* = 0.523, *p* < 0.001; G3 variance: *t* = 10.287, *d* = 0.824, *p* < 0.001). Furthermore, during movie watching, network positions shift closer to the center of the gradient compared to the resting state (see **STab.1** for more information on the network-level results). An animation of the observed gradient compression in the DyNAMiC dataset can be viewed in https://osf.io/vbxkw.

**Figure 4.**
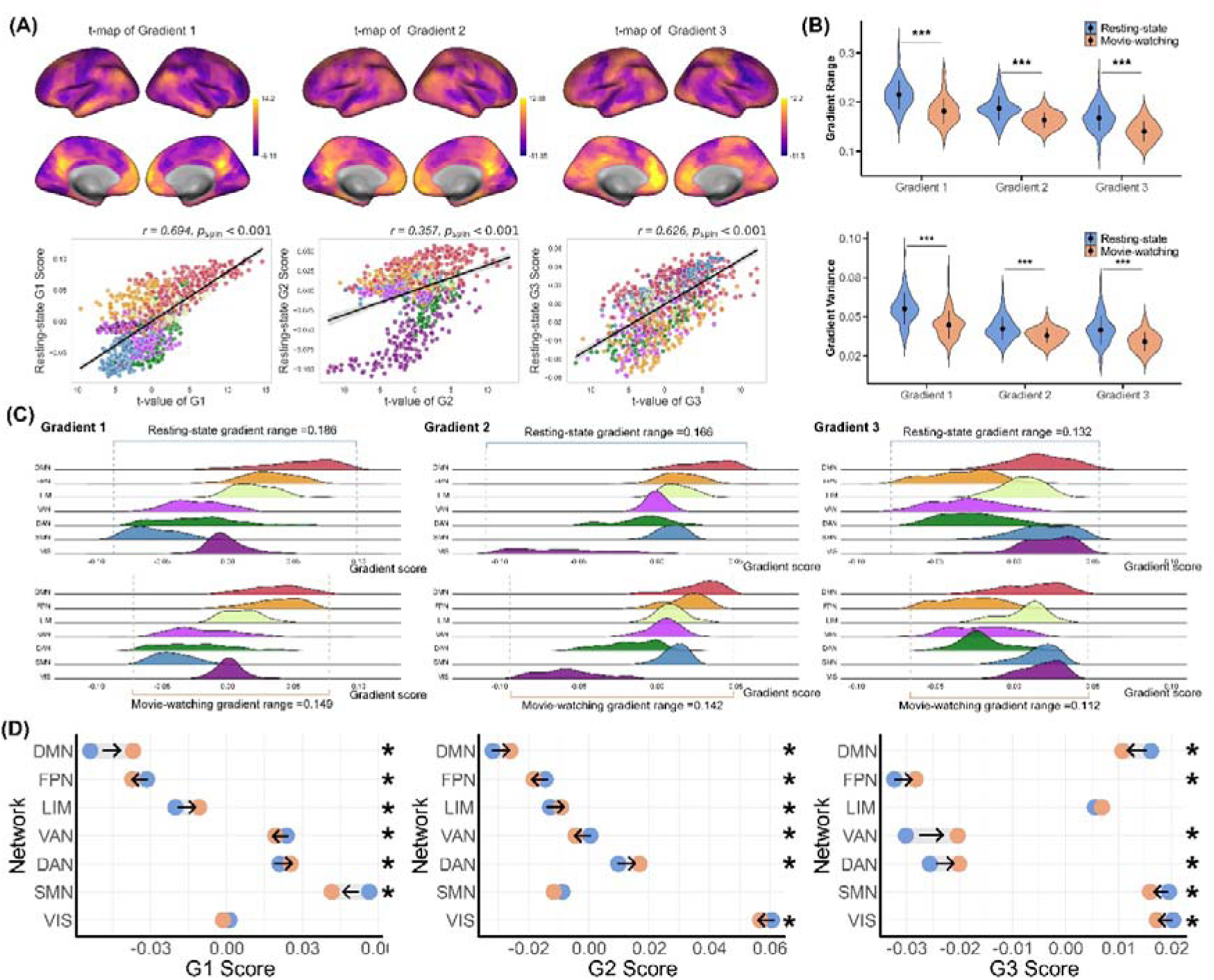
Regional, global, and network-level gradient differences between resting state and movie watching in DyNAMiC dataset. **(A)** The t-map of the difference in the regional gradient score between the two states, and the correlation between the t-map and resting-state gradient score. The significance of the correlation was estimated by using the spin permutation test. The shaded zone represents 95% confidence interval. **(B)** The violin plot illustrates that the gradient ranges and variances of the three gradients are higher during rest compared to naturalistic processing. **(C)** The distribution of the gradient score for Yeo’s 7 functional networks at two states. **(D)** The difference in network gradient score between the two states. The dot represents the network gradient score, and the arrow indicates the moving direction of the network position on the gradient axis when transitioning from rest to movie watching. VIS=visual network; SMN=somatomotor network; DAN=dorsal attention network; VAN=ventral attention network; LIM=limbic network; FPN=frontoparietal network; DMN= default mode network.

The overall results indicate a reliable compression pattern of the gradient during state transitions, implying that compared to resting state, naturalistic processing induces more homogenous connectivity profiles across regions, leading to a compressed functional hierarchy of the brain organization that was established in resting state.

### Functional integration mediates the state-driven gradient compression

To test whether the degree of gradient compression is mediated by functional integration and segregation, a mediation analysis was carried out between state-dependent differences in gradient range and PC and SS for each state. First, in Cam-CAN, higher PC was observed during movie-watching than resting-state (PC_Rest_ = 0.634 ± 0.044, PC_Movie_ = 0.683 ± 0.033, *t* = −20.120, *d* = −0.986, *p* < 0.001), while no significant state difference was revealed in functional segregation index (SS_Rest_ = 0.456 ± 0.084, SS_Movie_ = 0.454 ± 0.083, *t* = 0.426, *d* = 0.028, *p* = 0.670, **Fig.5A**). For both states, we found robust negative correlations between PC and gradient metrics of the V-S and S-A axis and positive correlations between SS and gradient metrics for all three gradients (**Fig.5B**). Given the significant state difference in the PC and the strong association between PC and gradient metrics, mediation models were constructed to test whether functional integration mediates the state-driven gradient alteration. The mediation analysis results indicate that PC partially mediates the alteration of gradient metrics due to state transitions, except for the variance of G3 (**Tab.3**).

**Figure 5.**
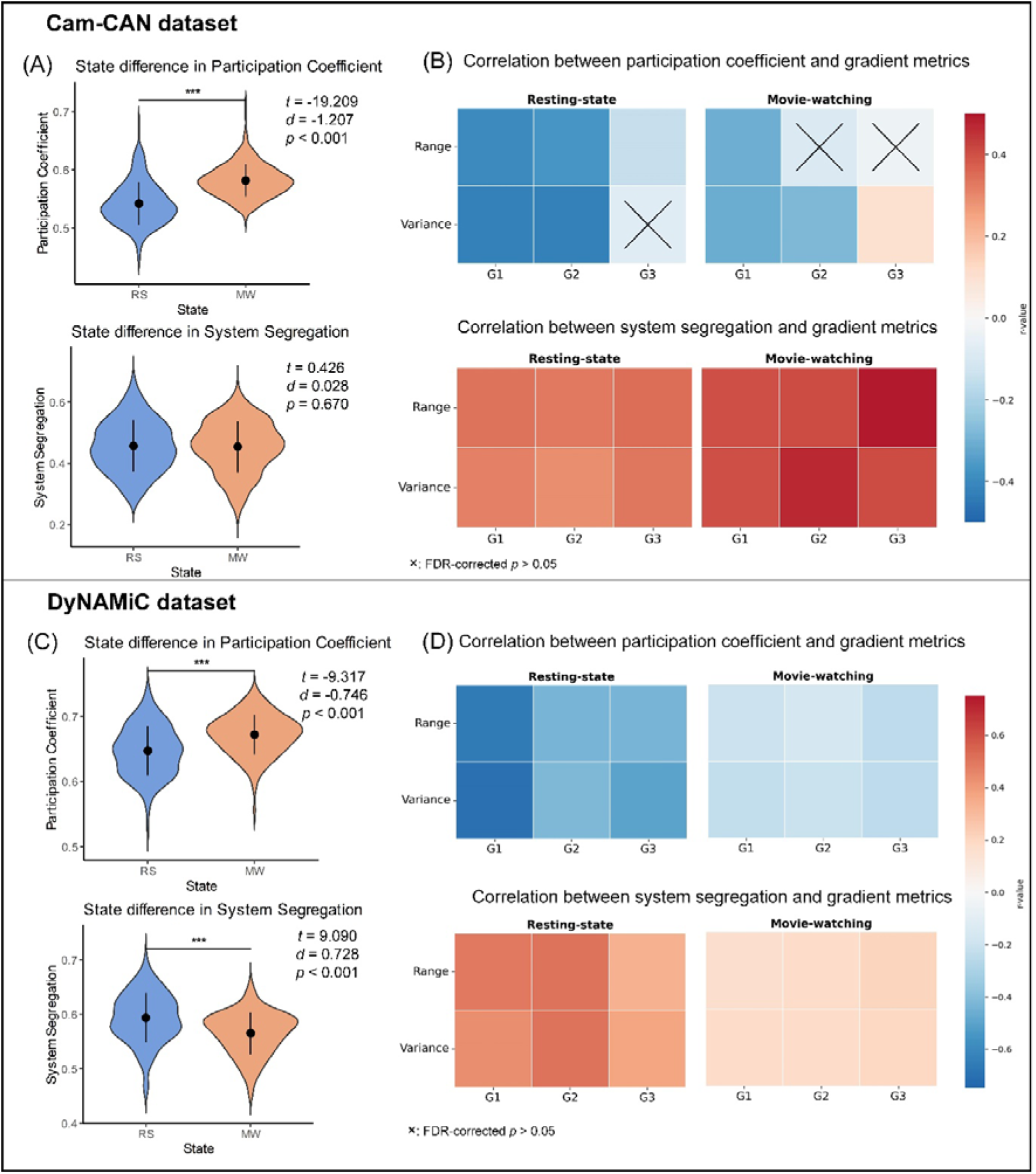
State difference in functional integration/segregation profiles and their associations with gradient metrics. **(A/C)** State difference in functional integration and segregation. The functional integration was estimated by participation coefficient and functional segregation was measured by system segregation. **(B/D)** The correlation between global gradient metrics and functional integration and segregation measures, with age and covariance controlled. RS=resting state; MW=movie watching. ***: *p*<0.001.

**Table 3.** Mediation effect of functional integration and segregation on state-driven alterations in global gradient metrics.

| Gradient Metrics | Est (Path a) | Est (Path b) | Est (Path ab) | Est (Path c') |
| --- | --- | --- | --- | --- |
| **Cam-CAN- Participation coefficient as mediator** |  |  |  |  |
| G1-Range | 0.359** | -0.250*** | 0.092* | -0.256*** |
| G2-Range | 0.359** | -0.470*** | 0.169** | -0.136 |
| G3-Range | 0.359** | -0.100 | -0.036 | 0.035 |
| G1-Variance | 0.359** | -0.371*** | -0.133** | -0.314** |
| G2- Variance | 0.359** | -0.351*** | -0.157** | -0.260** |
| G3- Variance | 0.359** | 0.005 | 0.002 | -0.272* |
| **DyNAMiC- Participation coefficient as mediator** |  |  |  |  |
| G1-Range | 0.348*** | -0.604*** | -0.210*** | -0.309*** |
| G2-Range | 0.348*** | -0.368*** | -0.128*** | -0.384*** |
| G3-Range | 0.348*** | -0.399*** | -0.139*** | -0.376*** |
| G1-Variance | 0.348*** | -0.657*** | -0.228** | -0.255*** |
| G2- Variance | 0.348*** | -0.402*** | -0.140*** | -0.197*** |
| G3- Variance | 0.348*** | -0.476*** | -0.292*** | -0.165*** |
| **DyNAMiC- System segregation as mediator** |  |  |  |  |
| G1-Range | -0.334*** | 0.466*** | -0.156*** | -0.364*** |
| G2-Range | -0.334*** | 0.438*** | -0.146*** | -0.365*** |
| G3-Range | -0.334*** | 0.323*** | -0.108*** | -0.407*** |
| G1-Variance | -0.334*** | 0.456*** | -0.152*** | -0.331*** |
| G2- Variance | -0.334*** | 0.502*** | -0.168*** | -0.169** |
| G3- Variance | -0.334*** | 0.367*** | -0.123*** | -0.335*** |

Note: \*: $p < 0.05$ ; \*\*: $p < 0.01$ ; \*\*\*: $p < 0.001$ .

In the DyNAMiC dataset, we similarly observed increased functional integration during movie when compared to rest (PC_Rest_ = 0.653 ± 0.040, PC_Movie_ = 0.671 ± 0.036, *t* = −9.318, *d* = −0.717, *p* < 0.001). However, in contrast to Cam-CAN, we also observed decreased functional segregation (SS_Rest_ = 0.594 ± 0.045, SS_Movie_ = 0.565 ± 0.039, *t* = 9.09, *d* = 0.728, *p* < 0.001, **Fig.5C**). Notably, consistent negative correlations were observed between PC and gradient metrics, in addition to positive correlations between SS and gradient metrics, across all three main gradients with age and covariance controlled (**Fig.5D**). Further mediation analyses revealed that the functional integration and segregation mediate the relationship between state and gradient metrics across main gradients (**Tab.3**).

Importantly, the above results largely persisted when PCs were calculated based on thresholded functional connectivity matrices at different sparsity thresholds (from 0.05 to 0.5 with the step of 0.05), especially those with higher sparsity levels that only take strongest connections into consideration (**SFig.4, STab.2**). These findings suggest that naturalistic processing enhances functional integration while reducing functional segregation, as reflected by increased interactions between functional networks, ultimately resulting in a more compressed functional cortical hierarchy in the brain.

### Aging amplifies the gradient compression from rest to movie

We next examined the hypothesis that older subjects may exhibit greater gradient compression when transitioning from rest to movie-watching as compared to their younger counterparts. The results from Cam-CAN showed that the state-age interaction significantly predicted all the gradient metrics (**Fig.6A**). Specifically, the state-age interaction showed significant positive associations with the range and variance of G1 (G1-range: *beta* = 0.135, *p* = 0.019; G1-variance: *beta* = 0.179, *p* = 0.005), and negatively predicted the range and variance of G2 and G3 (G2-range: *beta* = −0.203, *p* < 0.001; G2-variance: *beta* = −0.149, *p* = 0.013; G3-range: *beta* = −0.222, *p* < 0.001; G3-variance: *beta* = −0.188, *p* < 0.001, **Tab.4**). These results indicate that younger adults show greater gradient compression and reduced gradient variance along the V-S axis when transitioning from rest to movie-watching compared to older adults, while the opposite effect was observed for the S-A and insular-DMN axes. In other words, the S-A and insular-DMN axis showed greater state-driven gradient compression in older adults. The directionality of the observed state-age interaction effects is shown in the **Fig.6B**.

**Figure 6.**
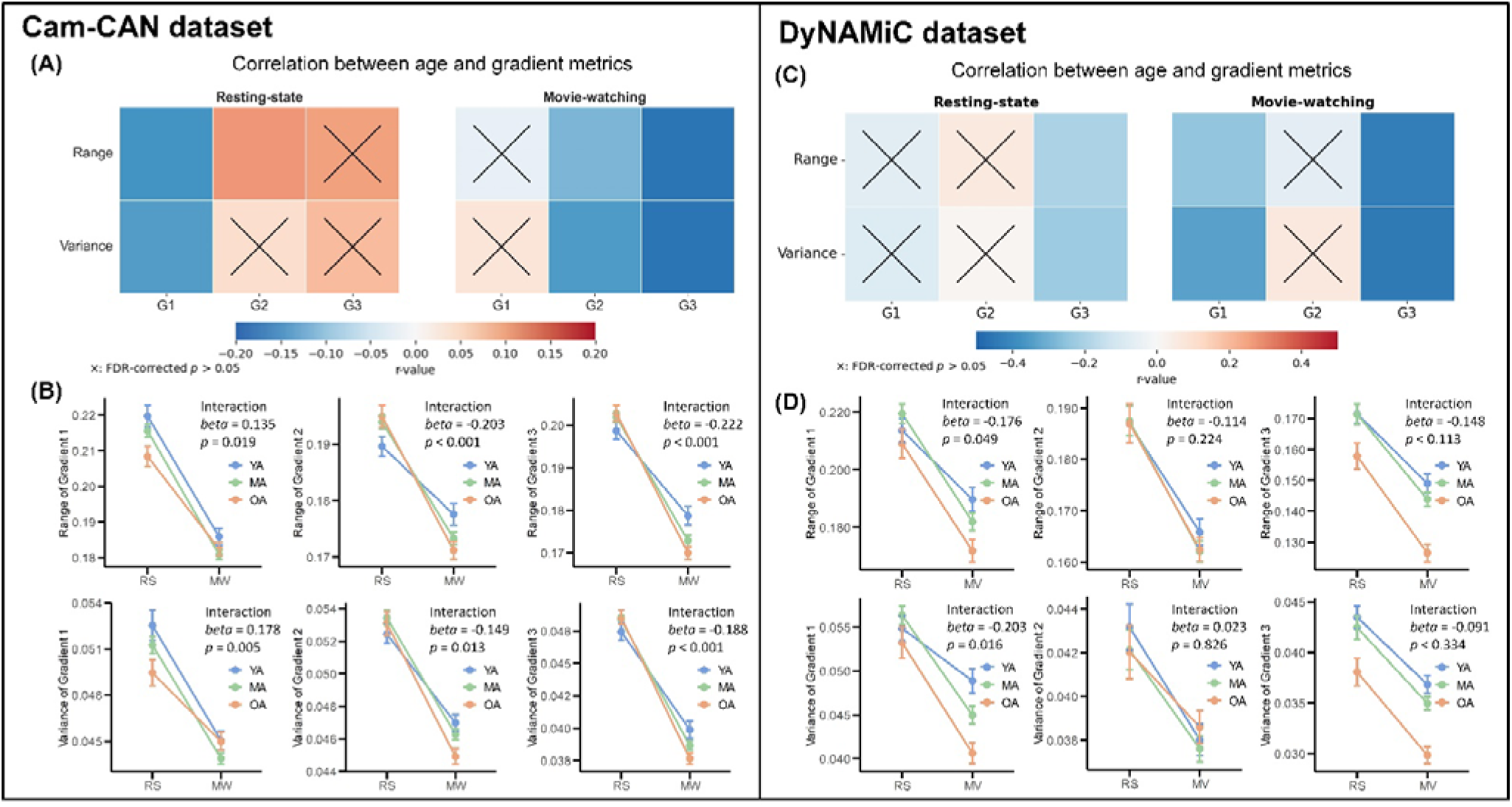
The moderation effect of age on the state-driven gradient alterations. **(A/C)** Correlation between age and gradient metrics. **(B/D)** The effect of age-state interaction on gradient metrics. For visualization the participants were assigned into three age groups: YA: age<35 years old; MA: age between 35 and 60 years old; OA: age>60 years old. RS=resting state; MW=movie watching; YA=younger adults’ group; MA= middle-aged group; OA=older adults’ group.

**Table 4.** Moderation effect of age on state-driven gradient alterations.

| Interaction | Gradient Metrics | *Beta* | *se* | *t-value* | *p-value* |
| --- | --- | --- | --- | --- | --- |
| **Cam-CAN** |  |  |  |  |  |
| State*Age | G1-Range | 0.135 | 0.058 | 2.341 | 0.019 |
| State*Age | G2-Range | -0.203 | 0.061 | -3.366 | <0.001 |
| State*Age | G3-Range | -0.222 | 0.054 | -4.096 | <0.001 |
| State*Age | G1-Std | 0.178 | 0.063 | 2.801 | 0.005 |
| State*Age | G2-Std | -0.149 | 0.060 | -2.500 | 0.013 |
| State*Age | G3-Std | -0.188 | 0.052 | -3.653 | <0.001 |
| **DyNAmiC** |  |  |  |  |  |
| State*Age | G1-Range | -0.176 | 0.089 | -1.976 | 0.049 |
| State*Age | G2-Range | -0.114 | 0.098 | -1.168 | 0.244 |
| State*Age | G3-Range | -0.148 | 0.092 | -1.592 | 0.113 |
| State*Age | G1-Std | -0.203 | 0.096 | -2.119 | 0.016 |
| State*Age | G2-Std | 0.023 | 0.107 | 0.219 | 0.826 |
| State*Age | G3-Std | -0.091 | 0.096 | -0.948 | 0.344 |

These results confirm our third hypothesis regarding the S-A and insular-DMN axes, substantiating that state-driven gradient compression is intensified with aging.

These conclusions were further supported by similar observations in DyNAMiC, where age was found to be negatively associated with the gradient range and variance of G3 (M-R axis) at rest, as well as G1 (S-A axis) and G3 during movie-watching (**Fig.6C**). Moderation analyses revealed a significant state-by-age interaction on G1 range (*beta* = −0.176, *p* = 0.049) and variance (*beta* = −0.203, *p* = 0.016), indicating that aging is associated with greater gradient compression when shifting from rest to movie watching, especially for the S-A axis (**Fig. 6D& Tab.4**).

### Gradient compression predicts cognitive performance

By constructing hierarchical multiple regression models, we tested whether the extent of gradient compression could predict cognitive performance. The overall results are illustrated in **Fig.7**. Specifically, across the full sample of the Cam-CAN dataset, we found the M2 with G2 compression added to the model could significantly predict emotional memory Recognition scores (*Adj.R^2^*= 0.450, Δ*R^2^* = 0.017, *F* = 3.020, *p* = 0.048) over and above base model M0 and M1 (*Adj.R^2^* = 0.438, Δ*R^2^* = 0.008, *F* = 1.438, *p* = 0.240), which failed to predict recognition score, and neither did M3 *(Adj.R^2^* = 0.447, Δ*R^2^* = 0.002, *F* = 0.446, *p* = 0.641). In the stepwise regression, ΔG2-range showed a significant negative predictive effect on recognition in both M2 (*beta* = −0.221, *t* =2.239, *p* = 0.026) and M3 (*beta* = −0.244, *t* =2.381, *p* = 0.018). However, no significant associations were found for other cognitive measures including Cattell, Priming, and Recollection. Furthermore, among older adults (age > 60 years), M2 predicted Cattell scores (*Adj.R^2^* = 0.236, Δ*R^2^* = 0.057, *F* = 4.219, *p* = 0.017). Within this group, only ΔG2-variance demonstrated a significant negative predictive effect on Cattell scores in both M2 (*beta* = −0.446, *t* =2.789, *p* = 0.006) and M3 (*beta* = −0.480, *t* =2.782, *p* = 0.006). No significant results were observed in younger or middle-aged groups.

**Figure 7.**
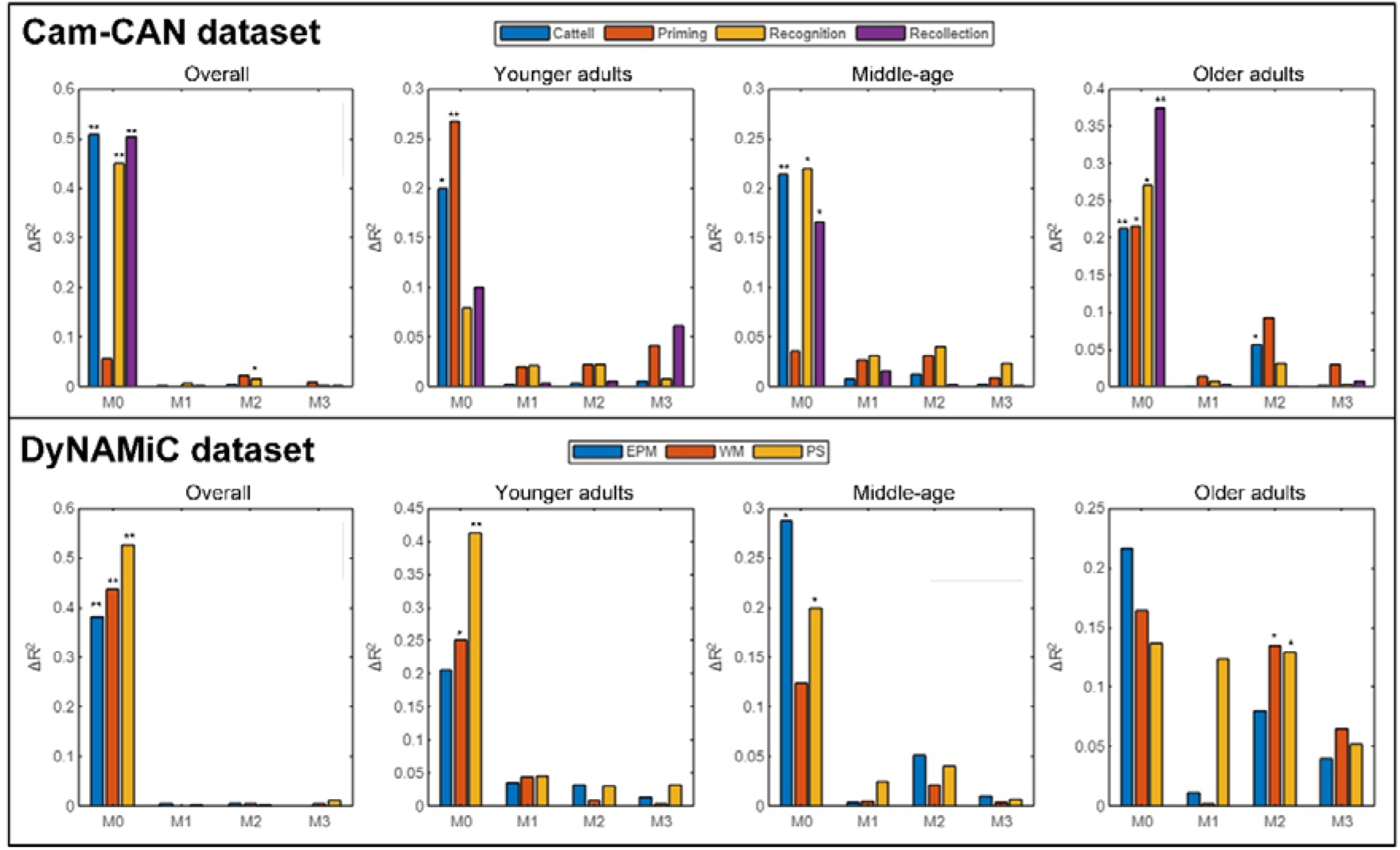
Predicting cognitive performance based on gradient compression. In Cam-CAN, gradient compression of G2 significantly predicted recognition scores across the overall sample, and it predicted Cattell scores in older adults. In DyNAMiC, gradient compression of G2 significantly predicted working memory and perceptual speed in older adults. The overall model was constructed across all age groups. EPM=episodic memory; WM=working memory; PS=perceptual speed. *: *p*<0.05 **: *p*<0.001.

In the overall DyNAMiC sample, no significant results were found across all cognitive performances when ΔG was entered into the regression model, nor in subsamples of younger and middle-aged adults. In older adults, however, we found that M2 (*Adj.R^2^* = 0.135, Δ*R^2^* = 0.135, *F* = 3.658, *p* = 0.035), which added G2 compression (ΔG2-range: *beta* = −0.827, *t* = −2.446, *p* = 0.019; ΔG2-variance: *beta* = −0.922, *t* = −2.705, *p* = 0.010), showed significant improvement from the M1(*Adj.R^2^*= 0.020, Δ*R^2^* = 0.002, *F* = 0.047, *p* = 0.954) in predicting working memory. M3 with G3 compression didn’t show significant improvement in predicting performance (*Adj.R^2^* = 0.171, Δ*R^2^* = 0.065, *F* = 1.829, *p* = 0.175), while ΔG2-range (*beta* = −0.825, *t* = −2.468, *p* = 0.018) and ΔG2-variance (*beta* = −0.884, *t* = −2.622, *p* = 0.013) remained significance in the model. Similar findings were also revealed in predicting perceptual speed. The M2 (*Adj.R^2^* = 0.390, Δ*R^2^* = 0.129, *F* = 3.912, *p* = 0.029) significantly predict perceptual speed, over and above base model M0 and M1 (*Adj.R^2^* = 0.261, Δ*R^2^* = 0.124, *F* = 3.055, *p* = 0.058), whereas only ΔG2-variance (*beta* = −0.828, *t* = −2.460, *p* = 0.019) was negatively associated with perceptual speed. M3 didn’t show significant improvement (*Adj.R^2^* = 0.251, Δ*R^2^* = 0.052, *F* = 1.629, *p* = 0.211), while ΔG2-variance (*beta* = −0.876, *t* = −2.638, *p* = 0.012) remained significant in the model.

Moreover, the association between age, cognitive performance and gradient compression were examined. Significant age-related cognitive decline was observed (Cam-CAN: Cattell: *r* = −0.638, *p* < 0.001; Priming: *r* = −0.605, *p* < 0.001; Recognition: *r* = −0.529, *p* < 0.001; Recollection: *r* = −0.605, *p* < 0.001; DyNAMiC: Working memory: *r* = −0.634, *p* < 0.001; Episodic memory: *r* = −0.517, *p* <0.001; Perpetual speed, *r* = −0.710, *p* < 0.001, **SFig.5**). Age was negatively correlated with ΔG1-range (*r* = - 0.112, *p* = 0.023) and ΔG1-variance (*r* = −0.147, *p* = 0.003), and positively correlated with ΔG2-range(*r* = 0.197, *p* < 0.001), ΔG2-variance(*r* = 0.157, *p* = 0.001), ΔG3-range (*r* = 0.210, *p* < 0.001), and ΔG3-variance (*r* = 0.189, *p* < 0.001) in Cam-CAN. However, no significant associations between age and gradient compression were found in DyNAMiC (all P’s > 0.05).

Furthermore, while several significant cognition-gradient compression associations were observed in Cam-CAN, all of these fell below statistical significance thresholds (i.e., 0.05) after age was further controlled in analysis (**STab.4**). No significant correlations were found between cognitive performance and gradient compression in DyNAMiC (**STab.3**).

## Discussion

The human brain integrates diverse sensory signals into a coherent understanding via the macroscale functional organization of the cortex (Bernhardt et al., 2022; Margulies et al., 2016). In this study, we leveraged two large datasets, Cam-CAN and DyNAMiC, to examine how macroscale brain organization changes during state transitions, particularly from rest to movie-watching. We identified a compression pattern of brain organization during this transition, driven by increased integration and decreased segregation of functional networks. This pattern reflects enhanced global interaction and reduced local specialization that supports the encoding of sensory perceptions into high-order cognition and emotion during naturalistic processing. Notably, aging intensified this gradient compression, particularly along the S-A axis, and this compression was associated with cognitive performance, especially in older adults. These findings provide the first evidence that macroscale brain organization compresses when transitioning from rest to naturalistic viewing, illustrating the age relevance and cognitive implications of state-driven gradient compression.

Efficient neural processing minimizes energy expenditure while maintaining computational capacity by dynamically adapting the topological configuration of brain networks to sensory inputs and cognitive demands(Bassett & Sporns, 2017; Bullmore & Sporns, 2012; Shine et al., 2019). Our results indicate that during the transition from rest to movie-watching, the brain’s principal gradients shift toward the center of the distribution, indicating a progressive hierarchical compression. This finding is consistent with prior reports of gradient compression during transitions from rest to cognitively demanding tasks (Ito & Murray, 2023; X. Wang et al., 2020; H. Zhang et al., 2022). Moreover, recent work has demonstrated that during rich cognitive processing, the brain exhibits more homogeneous neural activity with redundant patterns (Owen & Manning, 2024). At rest, the main gradients represent an intrinsic hierarchical architecture, uninfluenced by external tasks (Cole et al., 2014; Finn et al., 2017). In contrast, movie-watching elicits a less distinct hierarchical organization, reflecting a continuously dynamic state that mirrors real-life cognitive engagement. Given the fundamental role of these gradients in supporting high-order psychological processes(Jung et al., 2022; Mckeown et al., 2020), the observed compression may indicate an adaptive strategy to facilitate the integration of sensory information with cognitive and affective processing when confronted with dynamic stimuli.

Investigating the mechanisms underlying gradient compression revealed that increased functional integration underpins this shift. During movie-watching, both datasets showed greater functional integration compared to rest; however, reduced functional segregation was observed specifically in the DyNAMiC dataset. These findings align with previous evidence indicating that as task demands increase, functional networks become more integrated and less modular (Keller et al., 2023; Shine et al., 2019; Wainstein et al., 2021). Increased functional integration, measured by increased connectivity between networks, and decreased segregation, evident in weakened connectivity within networks, reflect a transition in the brain’s operational mode. This reorganization involves a decline of specialized processing in localized neural circuits in favor of a global configuration that facilitates distributed network collaboration to enhance network effectiveness and support complex cognition (Chen et al., 2017; Power et al., 2010; Shine et al., 2019). Therefore, the reconfiguration during movie-watching may represent an adaptive mechanism that optimizes resource allocation when individuals are exposed to external stimuli with high cognitive demand (Bolton et al., 2018; L. Song et al., 2023). Importantly, our study provides the first evidence that increased integration and decreased segregation mediate gradient compression during state transitions, reflecting a functional reorganization that rebalances local specialization and global communication to achieve efficient information transfer across states (Greene et al., 2023; Shine & Poldrack, 2018).

Age-related gradient compression was observed in both two datasets. Previous studies have linked aging with increased functional integration, reduced gradient variance, and increased cortical dedifferentiation during resting-state fMRI (Chan et al., 2014; Spreng & Turner, 2019; X. Wang et al., 2024). Similarly, studies focusing on subcortical areas (e.g., hippocampus and striatum) have shown age-related reductions in connectivity between subcortical and cortical areas (Korkki et al., 2025; Nordin et al., 2024). Moreover, a morphological study indicate that the cortical hierarchical structure becomes less distinct with age, suggesting a trend towards dedifferentiation (J. Li et al., 2024). Our study extend these observations to naturalistic processing by demonstrating that aging is associated with reduced gradient range and variance during movie watching. In other words, older adults exhibit greater connectivity homogeneity during naturalistic movie watching (Ferreira et al., 2016; Goh, 2011; Sala-Llonch et al., 2014).We observed that gradient compression becomes more pronounced with advancing age during transitions from rest to movie-watching, particularly along the S-A axis, which spans from primary sensorimotor to transmodal association areas and appears to be a key locus of age-related functional dedifferentiation (Meeker et al., 2020; Rieck et al., 2021). While gradient compression during state transitions (i.e., rest to movie-watching) might serve as an adaptive mechanism for optimizing resource allocation under cognitive demand, the aging appears disproportionately affected by this process. This dedifferentiation may reflect the diminished capacity of the aging brain to modulate neural activity in response to environmental demands, possibly due to declines in the flexibility of large-scale networks—a key mechanism for adapting to cognitive and sensory challenges (T. Li et al., 2023; H. Xia et al., 2024). Additionally, age-related gradient compression may be driven by declines in neurochemical systems, such as dopamine and acetylcholine, which are essential for neural plasticity and network flexibility (Karalija et al., 2024; Koen & Rugg, 2019; Mooraj et al., 2025). Structural changes including loss of while matter integrity and synaptic pruning, may further contribute to the observed age-related gradient compression, (Brito et al., 2023; Kirch & Gollo, 2021; Pedersen et al., 2021).

The relationship between gradient compression and cognitive performance further underscores its significance. The HMR analyses linked gradient compression to cognitive performance, particularly in older adults. This negative association is consistent with prior evidence that age-related dedifferentiation contributes to cognitive variability in aging populations (Meeker et al., 2020). Reduced modularity and increased functional integration have been associated with poorer performance in working memory, fluid intelligence, and executive function (Baniqued et al., 2018; Gu et al., 2022). These findings suggest that a diminished capacity to maintain distinct functional boundaries between networks impairs the efficient allocation of cognitive resources during demanding tasks (Rieck et al., 2021). In younger individuals, abundant cognitive resources and efficient neural modulation support an optimal balance between specificity and integration, allowing dynamic reorganization during tasks such as movie-watching, where integration information into a coherent narrative is prioritized over isolated processing (H. Song et al., 2021). In contrast, aging leads to decline in functional specificity and excessive integration, particularly during naturalistic processing. Consequently, in older adults, gradient compression no longer reflects an adaptive mechanism but becomes a marker of cognitive resource depletion and network dedifferentiation. Notably, previous evident suggests that older adults who maintain stronger functional specificity may resist excessive integration and maintain modularity despite limited cognitive resource (Liu et al., 2024; Stanford et al., 2022), resulting in better cognitive performance. This dissociation between gradient compression and cognition in younger versus older individuals aligns with the brain maintenance hypothesis (Nyberg et al., 2012) and highlights markers of brain reserve characterized by a well-balanced segregation-integration dynamic.

While the main findings were consistent across both the Cam-CAN and DyNAMiC datasets, some differences merit discussion. The primary gradients identified in the two datasets were not entirely consistent, despite widespread reporting of the S-A axis in previous studies (Margulies et al., 2016). Variations in the order of these gradients across samples may arise from differences in participant demographics and experimental protocols (Cross et al., 2021; De Rosa et al., 2024; Huo et al., 2022). Additionally, variations in resting-state paradigms and scan durations (e.g., an 8-minute eyes-closed scan in Cam-CAN versus a 12-minute eyes-open scan in DyNAMiC) could influence gradient estimation. Despite these differences, the robustness of our findings is supported by the consistent patterns observed across datasets, particularly regarding the relationship between gradient compression, age, and cognitive performance. The replication of key results across independent samples underscores the reliability of the core mechanisms underlying gradient reorganization and its cognitive implications.

Some limitations of the current study should be noted. First, the length, content, and narrative coherence of the movie stimulus significantly modulate brain activity during movie-watching (Hasson, 2008; Yang et al., 2023). However, the impact of these features on functional gradients was not fully explored in this study. Future studies should utilize a broader variety of movie types to validate our findings and further examine the influence of narrative on functional gradients. Second, the current study focused on cortical hierarchy, excluding subcortical regions from the analysis. Given the critical role of subcortical areas in regulating cognitive and emotional processes (Janacsek et al., 2022), future research should incorporate these regions to gain a more comprehensive understanding of hierarchical processing in the human brain. Third, by using resting-state gradients as the Procrustes alignment benchmark for movie-watching data, we prioritized cross-state comparability over capturing task-specific cortical hierarchy, potentially masking unique gradients specific to naturalistic processing.

In conclusion, this study provides the first multi-state perspective on age-related macroscale brain organization. Our analysis of two large datasets demonstrates that cortical gradients compress during transitions from rest to naturalistic processing, marked by increased integration and decreased segregation. This compression, which is intensified by aging, reflects excessive functional dedifferentiation that is associated with cognitive decline. These findings enhance our understanding of the dynamic reorganization of brain networks across states and underscore the importance of preserving hierarchical flexibility to mitigate cognitive decline in the aging population.

## Footnotes

- The authors declare no competing financial interests.
- We would like to thank all participants and the staff of the Cam-CAN and DyNAMiC project.

This study was suppoted by Kavli Foundation (grant number 47062019) and the Norwegian Research Council (Centers of Excellence scheme, project number 10399117).

- Data from the Cam-CAN project is publicly available at https://camcan-archive.mrc-cbu.cam.ac.uk/dataaccess/. Data from the DyNAMiC project are not publicly available due to ethical restrictions but can be made available upon reasonable request from the senior author (A.S.).
- Author contributions: S.Y., M.Z., and A.S. designed research; S.Y., J.M., X.L., R.B., and L.Z. analyzed data; S.Y wrote the first draft of the paper; S.Y., R.B., M.Z., and A.S. edited the paper; S.Y and M.Z. wrote the paper.

## Acknowledgments

We would like to thank all participants and the staff of the Cam-CAN and DyNAMiC project. This study was suppoted by Kavli Foundation (grant number 47062019) and the Norwegian Research Council (Centers of Excellence scheme, project number 10399117).

## Reference

Baniqued, P. L., Gallen, C. L., Voss, M. W., Burzynska, A. Z., Wong, C. N., Cooke, G. E., Duffy, K., Fanning, J., Ehlers, D. K., Salerno, E. A., Aguiñaga, S., McAuley, E., Kramer, A. F., & D’Esposito, M. (2018). Brain Network Modularity Predicts Exercise-Related Executive Function Gains in Older Adults. Frontiers in Aging Neuroscience, 9. 10.3389/fnagi.2017.00426

Bassett, D. S., & Sporns, O. (2017). Network neuroscience. Nature Neuroscience, 20(3), 353–364. 10.1038/nn.4502

Bernhardt, B. C., Smallwood, J., Keilholz, S., & Margulies, D. S. (2022). Gradients in brain organization. NeuroImage, 251, 118987. 10.1016/j.neuroimage.2022.118987

Bethlehem, R. A. I., Paquola, C., Seidlitz, J., Ronan, L., Bernhardt, B., Consortium, C.-C., & Tsvetanov, K. A. (2020). Dispersion of functional gradients across the adult lifespan. NeuroImage, 222, 117299. 10.1016/j.neuroimage.2020.117299

Bola, M., & Sabel, B. A. (2015). Dynamic reorganization of brain functional networks during cognition. NeuroImage, 114, 398–413. 10.1016/j.neuroimage.2015.03.057

Bolton, T. A. W., Jochaut, D., Giraud, A.-L., & Van De Ville, D. (2018). Brain dynamics in ASD during movie-watching show idiosyncratic functional integration and segregation. Human Brain Mapping, 39(6), 2391–2404. 10.1002/hbm.24009

Brito, D. V. C., Esteves, F., Rajado, A. T., Silva, N., Araújo, I., Bragança, J., Castelo-Branco, P., & Nóbrega, C. (2023). Assessing cognitive decline in the aging brain: Lessons from rodent and human studies. Npj Aging, 9(1), 1–11. 10.1038/s41514-023-00120-6

Brown, J. A., Lee, A. J., Pasquini, L., & Seeley, W. W. (2022). A dynamic gradient architecture generates brain activity states. NeuroImage, 261, 119526. 10.1016/j.neuroimage.2022.119526

Bullmore, E., & Sporns, O. (2012). The economy of brain network organization. Nature Reviews Neuroscience, 13(5), 336–349. 10.1038/nrn3214

Burt, J. B., Helmer, M., Shinn, M., Anticevic, A., & Murray, J. D. (2020). Generative modeling of brain maps with spatial autocorrelation. NeuroImage, 220, 117038. 10.1016/j.neuroimage.2020.117038

Chakraborty, S., Haast, R. A. M., Onuska, K. M., Kanel, P., Prado, M. A. M., Prado, V. F., Khan, A. R., & Schmitz, T. W. (2024). Multimodal gradients of basal forebrain connectivity across the neocortex. Nature Communications, 15(1), 8990. 10.1038/s41467-024-53148-x

Chan, M. Y., Park, D. C., Savalia, N. K., Petersen, S. E., & Wig, G. S. (2014). Decreased segregation of brain systems across the healthy adult lifespan. Proceedings of the National Academy of Sciences, 111(46), E4997–E5006. 10.1073/pnas.1415122111

Chang, L. J., Jolly, E., Cheong, J. H., Rapuano, K. M., Greenstein, N., Chen, P.-H. A., & Manning, J. R. (2021). Endogenous variation in ventromedial prefrontal cortex state dynamics during naturalistic viewing reflects affective experience. Science Advances, 7(17), eabf7129. 10.1126/sciadv.abf7129

Chen, Y., Wang, S., Hilgetag, C. C., & Zhou, C. (2017). Features of spatial and functional segregation and integration of the primate connectome revealed by trade-off between wiring cost and efficiency. PLOS Computational Biology, 13(9), e1005776. 10.1371/journal.pcbi.1005776

Coifman, R. R., Lafon, S., Lee, A. B., Maggioni, M., Nadler, B., Warner, F., & Zucker, S. W. (2005). Geometric diffusions as a tool for harmonic analysis and structure definition of data: Diffusion maps. Proceedings of the National Academy of Sciences, 102(21), 7426–7431. 10.1073/pnas.0500334102

Cole, M. W., Bassett, D. S., Power, J. D., Braver, T. S., & Petersen, S. E. (2014). Intrinsic and Task-Evoked Network Architectures of the Human Brain. Neuron, 83(1), 238–251. 10.1016/j.neuron.2014.05.014

Dong, H.-M., Zhang, X.-H., Labache, L., Zhang, S., Ooi, L. Q. R., Yeo, B. T. T., Margulies, D. S., Holmes, A. J., & Zuo, X.-N. (2024). Ventral attention network connectivity is linked to cortical maturation and cognitive ability in childhood. Nature Neuroscience, 1–12. 10.1038/s41593-024-01736-x

Esmaeili, M., Bjørkeli, E. B., Pedersen, R., Falahati, F., Johansson, J., Nordin, K., Karalija, N., Bäckman, L., Nyberg, L., & Salami, A. (2025). Brain-Cognitive Gaps in relation to Dopamine and Health-related Factors: Insights from AI-Driven Functional Connectome Predictions. eLife, 14. 10.7554/eLife.104053.1

Ferreira, L. K., Regina, A. C. B., Kovacevic, N., Martin, M. da G. M., Santos, P. P., Carneiro, C. de G., Kerr, D. S., Amaro, E., Jr, McIntosh, A. R., & Busatto, G. F. (2016). Aging Effects on Whole-Brain Functional Connectivity in Adults Free of Cognitive and Psychiatric Disorders. Cerebral Cortex, 26(9), 3851–3865. 10.1093/cercor/bhv190

Finn, E. S. (2021). Is it time to put rest to rest? Trends in Cognitive Sciences, 25(12), 1021–1032. 10.1016/j.tics.2021.09.005

Finn, E. S., & Bandettini, P. A. (2021). Movie-watching outperforms rest for functional connectivity-based prediction of behavior. NeuroImage, 235, 117963. 10.1016/j.neuroimage.2021.117963

Finn, E. S., Scheinost, D., Finn, D. M., Shen, X., Papademetris, X., & Constable, R. T. (2017). Can brain state be manipulated to emphasize individual differences in functional connectivity? NeuroImage, 160, 140–151. 10.1016/j.neuroimage.2017.03.064

Gale, D. J., Areshenkoff, C. N., Standage, D. I., Nashed, J. Y., Markello, R. D., Flanagan, J. R., Smallwood, J., & Gallivan, J. P. (2022). Distinct patterns of cortical manifold expansion and contraction underlie human sensorimotor adaptation. Proceedings of the National Academy of Sciences, 119(52), e2209960119. 10.1073/pnas.2209960119

Gaser, C., Dahnke, R., Thompson, P. M., Kurth, F., Luders, E., & the Alzheimer’s Disease Neuroimaging Initiative. (2024). CAT: A computational anatomy toolbox for the analysis of structural MRI data. GigaScience, 13, giae049. 10.1093/gigascience/giae049

Girn, M., Roseman, L., Bernhardt, B., Smallwood, J., Carhart-Harris, R., & Nathan Spreng, R. (2022). Serotonergic psychedelic drugs LSD and psilocybin reduce the hierarchical differentiation of unimodal and transmodal cortex. NeuroImage, 256, 119220. 10.1016/j.neuroimage.2022.119220

Goh, J. O. S. (2011). Functional Dedifferentiation and Altered Connectivity in Older Adults: Neural Accounts of Cognitive Aging. Aging and Disease, 2(1), 30–48.

Greene, A. S., Horien, C., Barson, D., Scheinost, D., & Constable, R. T. (2023). Why is everyone talking about brain state? Trends in Neurosciences, 46(7), 508–524. 10.1016/j.tins.2023.04.001

Gu, Y., Li, L., Zhang, Y., Ma, J., Yang, C., Xiao, Y., Shu, N., Can, C., Lin, Y., & Dai, Z. (2022). The overlapping modular organization of human brain functional networks across the adult lifespan. NeuroImage, 253, 119125. 10.1016/j.neuroimage.2022.119125

Hasson, U. (2008). Neurocinematics: The Neuroscience of Film. Projections, 2, 1–26.

Huang, Z., Mashour, G. A., & Hudetz, A. G. (2023). Functional geometry of the cortex encodes dimensions of consciousness. Nature Communications, 14(1), 72. 10.1038/s41467-022-35764-7

Huntenburg, J. M., Bazin, P.-L., & Margulies, D. S. (2018). Large-Scale Gradients in Human Cortical Organization. Trends in Cognitive Sciences, 22(1), 21–31. 10.1016/j.tics.2017.11.002

Ito, T., & Murray, J. D. (2023). Multitask representations in the human cortex transform along a sensory-to-motor hierarchy. Nature Neuroscience, 26(2), 306–315. 10.1038/s41593-022-01224-0

Janacsek, K., Evans, T. M., Kiss, M., Shah, L., Blumenfeld, H., & Ullman, M. T. (2022). Subcortical Cognition: The Fruit Below the Rind. Annual Review of Neuroscience, 45(Volume 45, 2022), 361–386. 10.1146/annurev-neuro-110920-013544

Johansson, J., Nordin, K., Pedersen, R., Karalija, N., Papenberg, G., Andersson, M., Korkki, S. M., Riklund, K., Guitart-Masip, M., Rieckmann, A., Bäckman, L., Nyberg, L., & Salami, A. (2023). Biphasic patterns of age-related differences in dopamine D1 receptors across the adult lifespan. Cell Reports, 42(9). 10.1016/j.celrep.2023.113107

Jung, H., Wager, T. D., & Carter, R. M. (2022). Novel Cognitive Functions Arise at the Convergence of Macroscale Gradients. Journal of Cognitive Neuroscience, 34(3), 381–396. 10.1162/jocn_a_01803

Karalija, N., Papenberg, G., Johansson, J., Wåhlin, A., Salami, A., Andersson, M., Axelsson, J., Kuznetsov, D., Riklund, K., Lövdén, M., Lindenberger, U., Bäckman, L., & Nyberg, L. (2024). Longitudinal support for the correlative triad among aging, dopamine D2-like receptor loss, and memory decline. Neurobiology of Aging, 136, 125–132. 10.1016/j.neurobiolaging.2024.02.001

Katsumi, Y., Theriault, J. E., Quigley, K. S., & Barrett, L. F. (2022). Allostasis as a core feature of hierarchical gradients in the human brain. Network Neuroscience, 6(4), 1010–1031. 10.1162/netn_a_00240

Katsumi, Y., Zhang, J., Chen, D., Kamona, N., Bunce, J. G., Hutchinson, J. B., Yarossi, M., Tunik, E., Dickerson, B. C., Quigley, K. S., & Barrett, L. F. (2023). Correspondence of functional connectivity gradients across human isocortex, cerebellum, and hippocampus. Communications Biology, 6(1), 1–13. 10.1038/s42003-023-04796-0

Keller, A. S., Sydnor, V. J., Pines, A., Fair, D. A., Bassett, D. S., & Satterthwaite, T. D. (2023). Hierarchical functional system development supports executive function. Trends in Cognitive Sciences, 27(2), 160–174. 10.1016/j.tics.2022.11.005

Kirch, C., & Gollo, L. L. (2021). Single-neuron dynamical effects of dendritic pruning implicated in aging and neurodegeneration: Towards a measure of neuronal reserve. Scientific Reports, 11(1), 1309. 10.1038/s41598-020-78815-z

Knodt, A. R., Elliott, M. L., Whitman, E. T., Winn, A., Addae, A., Ireland, D., Poulton, R., Ramrakha, S., Caspi, A., Moffitt, T. E., & Hariri, A. R. (2023). Test–retest reliability and predictive utility of a macroscale principal functional connectivity gradient. Human Brain Mapping, 44(18), 6399– 6417. 10.1002/hbm.26517

Koen, J. D., & Rugg, M. D. (2019). Neural Dedifferentiation in the Aging Brain. Trends in Cognitive Sciences, 23(7), 547–559. 10.1016/j.tics.2019.04.012

Korkki, S. M., Johansson, J., Nordin, K., Pedersen, R., Bäckman, L., Rieckmann, A., & Salami, A. (2025). Dedifferentiation of caudate functional organization is linked to reduced D1 dopamine receptor availability and poorer memory function in aging. Imaging Neuroscience, 3, imag_a_00462. 10.1162/imag_a_00462

Langs, G., Golland, P., & Ghosh, S. S. (2015). Predicting Activation Across Individuals with Resting-State Functional Connectivity Based Multi-Atlas Label Fusion. Medical Image Computing and Computer-Assisted Intervention⍰: MICCAI… International Conference on Medical Image Computing and Computer-Assisted Intervention, 9350, 313. 10.1007/978-3-319-24571-3_38

Lei, W., Xiao, Q., Wang, C., Cai, Z., Lu, G., Su, L., & Zhong, Y. (2023). The disruption of functional connectome gradient revealing networks imbalance in pediatric bipolar disorder. Journal of Psychiatric Research, 164, 72–79. 10.1016/j.jpsychires.2023.05.084

Li, J., Zhang, C., Meng, Y., Yang, S., Xia, J., Chen, H., & Liao, W. (2024). Morphometric brain organization across the human lifespan reveals increased dispersion linked to cognitive performance. PLOS Biology, 22(6), e3002647. 10.1371/journal.pbio.3002647

Li, T., Xia, H., Li, H., He, Q., & Chen, A. (2023). Functional Connectivity Alterations of Cognitive Flexibility in Aging: Different Patterns of Global and Local Switch Costs. The Journals of Gerontology: Series B, 78(10), 1651–1658. 10.1093/geronb/gbad092

Liu, H., Jing, J., Jiang, J., Wen, W., Zhu, W., Li, Z., Pan, Y., Cai, X., Liu, C., Zhou, Y., Meng, X., Wang, Y., Li, H., Jiang, Y., Zheng, H., Wang, S., Niu, H., Kochan, N., Brodaty, H.,… Wang, Y. (2024). Exploring the link between brain topological resilience and cognitive performance in the context of aging and vascular risk factors: A cross-ethnicity population-based study. Science Bulletin, 69(17), 2735–2744. 10.1016/j.scib.2024.04.018

Margulies, D. S., Ghosh, S. S., Goulas, A., Falkiewicz, M., Huntenburg, J. M., Langs, G., Bezgin, G., Eickhoff, S. B., Castellanos, F. X., Petrides, M., Jefferies, E., & Smallwood, J. (2016). Situating the default-mode network along a principal gradient of macroscale cortical organization. Proceedings of the National Academy of Sciences, 113(44), 12574–12579. 10.1073/pnas.1608282113

Mckeown, B., Strawson, W. H., Wang, H.-T., Karapanagiotidis, T., Vos de Wael, R., Benkarim, O., Turnbull, A., Margulies, D., Jefferies, E., McCall, C., Bernhardt, B., & Smallwood, J. (2020). The relationship between individual variation in macroscale functional gradients and distinct aspects of ongoing thought. NeuroImage, 220, 117072. 10.1016/j.neuroimage.2020.117072

Meeker, K. L., Ances, B. M., Gordon, B. A., Morris, J. C., Benzinger, T. L. S., & Waring, J. (2020). Default mode network dedifferentiation predicts cognitive performance in Alzheimer disease. Alzheimer’s & Dementia, 16(S4), e044790. 10.1002/alz.044790

Meer, J. N. van der, Breakspear, M., Chang, L. J., Sonkusare, S., & Cocchi, L. (2020). Movie viewing elicits rich and reliable brain state dynamics. Nat. Commun, 11, 5004.

Mooraj, Z., Salami, A., Campbell, K. L., Dahl, M. J., Kosciessa, J. Q., Nassar, M. R., Werkle-Bergner, M., Craik, F. I. M., Lindenberger, U., Mayr, U., Rajah, M. N., Raz, N., Nyberg, L., & Garrett, D. D. (2025). Toward a functional future for the cognitive neuroscience of human aging. Neuron, 113(1), 154–183. 10.1016/j.neuron.2024.12.008

Nordin, K., Gorbach, T., Pedersen, R., Panes Lundmark, V., Johansson, J., Andersson, M., McNulty, C., Riklund, K., Wåhlin, A., Papenberg, G., Kalpouzos, G., Bäckman, L., & Salami, A. (2022). DyNAMiC: A prospective longitudinal study of dopamine and brain connectomes: A new window into cognitive aging. Journal of Neuroscience Research, 100(6), 1296–1320. 10.1002/jnr.25039

Nordin, K., Pedersen, R., Falahati, F., Johansson, J., Grill, F., Andersson, M., Korkki, S. M., Bäckman, L., Zalesky, A., Rieckmann, A., Nyberg, L., & Salami, A. (2024). Two long-axis dimensions of hippocampal-cortical integration support memory function across the adult lifespan. eLife, 13. 10.7554/eLife.97658.1

Nyberg, L., Lövdén, M., Riklund, K., Lindenberger, U., & Bäckman, L. (2012). Memory aging and brain maintenance. Trends in Cognitive Sciences, 16(5), 292–305. 10.1016/j.tics.2012.04.005

Ottoy, J., Kang, M. S., Tan, J. X. M., Boone, L., Vos de Wael, R., Park, B., Bezgin, G., Lussier, F. Z., Pascoal, T. A., Rahmouni, N., Stevenson, J., Fernandez Arias, J., Therriault, J., Hong, S.-J., Stefanovic, B., McLaurin, J., Soucy, J.-P., Gauthier, S., Bernhardt, B. C.,… Goubran, M. (2024). Tau follows principal axes of functional and structural brain organization in Alzheimer’s disease. Nature Communications, 15(1), 5031. 10.1038/s41467-024-49300-2

Owen, L. L. W., & Manning, J. R. (2024). High-level cognition is supported by information-rich but compressible brain activity patterns. Proceedings of the National Academy of Sciences, 121(35), e2400082121. 10.1073/pnas.2400082121

Pedersen, R., Geerligs, L., Andersson, M., Gorbach, T., Avelar-Pereira, B., Wåhlin, A., Rieckmann, A., Nyberg, L., & Salami, A. (2021). When functional blurring becomes deleterious: Reduced system segregation is associated with less white matter integrity and cognitive decline in aging. NeuroImage, 242, 118449. 10.1016/j.neuroimage.2021.118449

Pedersen, R., Johansson, J., & Salami, A. (2023). Dopamine D1-signaling modulates maintenance of functional network segregation in aging. Aging Brain, 3, 100079. 10.1016/j.nbas.2023.100079

Pines, A. R., Larsen, B., Cui, Z., Sydnor, V. J., Bertolero, M. A., Adebimpe, A., Alexander-Bloch, A. F., Davatzikos, C., Fair, D. A., Gur, R. C., Gur, R. E., Li, H., Milham, M. P., Moore, T. M., Murtha, K., Parkes, L., Thompson-Schill, S. L., Shanmugan, S., Shinohara, R. T.,… Satterthwaite, T. D. (2022). Dissociable multi-scale patterns of development in personalized brain networks. Nature Communications, 13(1), 2647. 10.1038/s41467-022-30244-4

Power, J. D., Barnes, K. A., Snyder, A. Z., Schlaggar, B. L., & Petersen, S. E. (2012). Spurious but systematic correlations in functional connectivity MRI networks arise from subject motion. NeuroImage, 59(3), 2142–2154. 10.1016/j.neuroimage.2011.10.018

Power, J. D., Fair, D. A., Schlaggar, B. L., & Petersen, S. E. (2010). The Development of Human Functional Brain Networks. Neuron, 67(5), 735–748. 10.1016/j.neuron.2010.08.017

Power, J. D., Schlaggar, B. L., Lessov-Schlaggar, C. N., & Petersen, S. E. (2013). Evidence for Hubs in Human Functional Brain Networks. Neuron, 79(4), 798–813. 10.1016/j.neuron.2013.07.035

Ren, Y., Nguyen, V. T., Sonkusare, S., Lv, J., Pang, T., Guo, L., Eickhoff, S. B., Breakspear, M., & Guo, C. C. (2018). Effective connectivity of the anterior hippocampus predicts recollection confidence during natural memory retrieval. Nature Communications, 9(1), 4875. 10.1038/s41467-018-07325-4

Rieck, J. R., Baracchini, G., Nichol, D., Abdi, H., & Grady, C. L. (2021). Reconfiguration and dedifferentiation of functional networks during cognitive control across the adult lifespan. Neurobiology of Aging, 106, 80–94. 10.1016/j.neurobiolaging.2021.03.019

Sala-Llonch, R., Junqué, C., Arenaza-Urquijo, E. M., Vidal-Piñeiro, D., Valls-Pedret, C., Palacios, E. M., Domènech, S., Salvà, A., Bargalló, N., & Bartrés-Faz, D. (2014). Changes in whole-brain functional networks and memory performance in aging. Neurobiology of Aging, 35(10), 2193–2202. 10.1016/j.neurobiolaging.2014.04.007

Samara, A., Eilbott, J., Margulies, D. S., Xu, T., & Vanderwal, T. (2023). Cortical gradients during naturalistic processing are hierarchical and modality-specific. NeuroImage, 271, 120023. 10.1016/j.neuroimage.2023.120023

Schaefer, A., Kong, R., Gordon, E. M., Laumann, T. O., Zuo, X.-N., Holmes, A. J., Eickhoff, S. B., & Yeo, B. T. T. (2018). Local-Global Parcellation of the Human Cerebral Cortex from Intrinsic Functional Connectivity MRI. Cerebral Cortex, 28(9), 3095–3114. 10.1093/cercor/bhx179

Seguin, C., Sporns, O., & Zalesky, A. (2023). Brain network communication: Concepts, models and applications. Nature Reviews Neuroscience, 24(9), 557–574. 10.1038/s41583-023-00718-5

Senkowski, D., & Engel, A. K. (2024). Multi-timescale neural dynamics for multisensory integration. Nature Reviews Neuroscience, 25(9), 625–642. 10.1038/s41583-024-00845-7

Shafto, M. A., Tyler, L. K., Dixon, M., Taylor, J. R., Rowe, J. B., Cusack, R., Calder, A. J., Marslen-Wilson, W. D., Duncan, J., Dalgleish, T., Henson, R. N., Brayne, C., Matthews, F. E., & Cam-CAN. (2014). The Cambridge Centre for Ageing and Neuroscience (Cam-CAN) study protocol: A cross-sectional, lifespan, multidisciplinary examination of healthy cognitive ageing. BMC Neurology, 14(1), 204. 10.1186/s12883-014-0204-1

Shine, J. M., Breakspear, M., Bell, P. T., Ehgoetz Martens, K. A., Shine, R., Koyejo, O., Sporns, O., & Poldrack, R. A. (2019). Human cognition involves the dynamic integration of neural activity and neuromodulatory systems. Nature Neuroscience, 22(2), 289–296. 10.1038/s41593-018-0312-0

Shine, J. M., & Poldrack, R. A. (2018). Principles of dynamic network reconfiguration across diverse brain states. NeuroImage, 180, 396–405. 10.1016/j.neuroimage.2017.08.010

Song, H., Park, B., Park, H., & Shim, W. M. (2021). Cognitive and Neural State Dynamics of Narrative Comprehension. Journal of Neuroscience, 41(43), 8972–8990. 10.1523/JNEUROSCI.0037-21.2021

Song, L., Ren, Y., Wang, K., Hou, Y., Nie, J., & He, X. (2023). Mapping the time-varying functional brain networks in response to naturalistic movie stimuli. Frontiers in Neuroscience, 17. 10.3389/fnins.2023.1199150

Spreng, R. N., & Turner, G. R. (2019). The Shifting Architecture of Cognition and Brain Function in Older Adulthood. Perspectives on Psychological Science, 14(4), 523–542. 10.1177/1745691619827511

Stanford, W. C., Mucha, P. J., & Dayan, E. (2022). A robust core architecture of functional brain networks supports topological resilience and cognitive performance in middle- and old-aged adults. Proceedings of the National Academy of Sciences, 119(44), e2203682119. 10.1073/pnas.2203682119

Su, K., Huang, Z., Li, Q., Fan, M., Li, T., & Yin, D. (2024). Dissociable functional responses along the posterior-anterior gradient of the frontal and parietal cortices revealed by parametric working memory and training. Brain Structure and Function, 229(7), 1681–1696. 10.1007/s00429-024-02834-z

Taylor, J. R., Williams, N., Cusack, R., Auer, T., Shafto, M. A., Dixon, M., Tyler, L. K., Cam-CAN, & Henson, R. N. (2017). The Cambridge Centre for Ageing and Neuroscience (Cam-CAN) data repository: Structural and functional MRI, MEG, and cognitive data from a cross-sectional adult lifespan sample. NeuroImage, 144, 262–269. 10.1016/j.neuroimage.2015.09.018

Tong, C., Liu, C., Zhang, K., Bo, B., Xia, Y., Yang, H., Feng, Y., & Liang, Z. (2022). Multimodal analysis demonstrating the shaping of functional gradients in the marmoset brain. Nature Communications, 13(1), 6584. 10.1038/s41467-022-34371-w

Tsvetanov, K. A., Henson, R. N. A., Tyler, L. K., Razi, A., Geerligs, L., Ham, T. E., Rowe, J. B., & Neuroscience, C. C. for A. and. (2016). Extrinsic and Intrinsic Brain Network Connectivity Maintains Cognition across the Lifespan Despite Accelerated Decay of Regional Brain Activation. Journal of Neuroscience, 36(11), 3115–3126. 10.1523/JNEUROSCI.2733-15.2016

Tsvetanov, K. A., Ye, Z., Hughes, L., Samu, D., Treder, M. S., Wolpe, N., Tyler, L. K., Rowe, J. B., & Neuroscience, for the C. C. for A. and. (2018). Activity and Connectivity Differences Underlying Inhibitory Control Across the Adult Life Span. Journal of Neuroscience, 38(36), 7887–7900. 10.1523/JNEUROSCI.2919-17.2018

Valk, S. L., Xu, T., Paquola, C., Park, B., Bethlehem, R. A. I., Vos de Wael, R., Royer, J., Masouleh, S. K., Bayrak, Ş., Kochunov, P., Yeo, B. T. T., Margulies, D., Smallwood, J., Eickhoff, S. B., & Bernhardt, B. C. (2022). Genetic and phylogenetic uncoupling of structure and function in human transmodal cortex. Nature Communications, 13(1), 2341. 10.1038/s41467-022-29886-1

Vatansever, D., Menon, D. K., Manktelow, A. E., Sahakian, B. J., & Stamatakis, E. A. (2015). Default Mode Dynamics for Global Functional Integration. Journal of Neuroscience, 35(46), 15254– 15262. 10.1523/JNEUROSCI.2135-15.2015

Vos de Wael, R., Benkarim, O., Paquola, C., Lariviere, S., Royer, J., Tavakol, S., Xu, T., Hong, S.-J., Langs, G., Valk, S., Misic, B., Milham, M., Margulies, D., Smallwood, J., & Bernhardt, B. C. (2020). BrainSpace: A toolbox for the analysis of macroscale gradients in neuroimaging and connectomics datasets. Communications Biology, 3(1), 1–10. 10.1038/s42003-020-0794-7

Wainstein, G., Rojas-Líbano, D., Medel, V., Alnæs, D., Kolskår, K. K., Endestad, T., Laeng, B., Ossandon, T., Crossley, N., Matar, E., & Shine, J. M. (2021). The ascending arousal system promotes optimal performance through mesoscale network integration in a visuospatial attentional task. Network Neuroscience, 5(4), 890–910. 10.1162/netn_a_00205

Wang, J., Ren, Y., Hu, X., Nguyen, V. T., Guo, L., Han, J., & Guo, C. C. (2017). Test–retest reliability of functional connectivity networks during naturalistic fMRI paradigms. Human Brain Mapping, 38(4), 2226–2241. 10.1002/hbm.23517

Wang, R., Liu, M., Cheng, X., Wu, Y., Hildebrandt, A., & Zhou, C. (2021). Segregation, integration, and balance of large-scale resting brain networks configure different cognitive abilities. Proceedings of the National Academy of Sciences, 118(23), e2022288118. 10.1073/pnas.2022288118

Wang, X., Huang, C.-C., Tsai, S.-J., Lin, C.-P., & Cai, Q. (2024). The aging trajectories of brain functional hierarchy and its impact on cognition across the adult lifespan. Frontiers in Aging Neuroscience, 16. 10.3389/fnagi.2024.1331574

Wang, X., Margulies, D. S., Smallwood, J., & Jefferies, E. (2020). A gradient from long-term memory to novel cognition: Transitions through default mode and executive cortex. NeuroImage, 220, 117074. 10.1016/j.neuroimage.2020.117074

Wu, W., & Hoffman, P. (2024). Functional integration and segregation during semantic cognition: Evidence across age groups. Cortex, 178, 157–173. 10.1016/j.cortex.2024.06.015

Xia, H., Hou, Y., Li, Q., & Chen, A. (2024). A meta-analysis of cognitive flexibility in aging: Perspective from functional network and lateralization. Human Brain Mapping, 45(14), e70031. 10.1002/hbm.70031

Xia, Y., Xia, M., Liu, J., Liao, X., Lei, T., Liang, X., Zhao, T., Shi, Z., Sun, L., Chen, X., Men, W., Wang, Y., Pan, Z., Luo, J., Peng, S., Chen, M., Hao, L., Tan, S., Gao, J.-H.,… He, Y. (2022). Development of functional connectome gradients during childhood and adolescence. Science Bulletin, 67(10), 1049–1061. 10.1016/j.scib.2022.01.002

Yang, E., Milisav, F., Kopal, J., Holmes, A. J., Mitsis, G. D., Misic, B., Finn, E. S., & Bzdok, D. (2023). The default network dominates neural responses to evolving movie stories. Nature Communications, 14(1), 4197. 10.1038/s41467-023-39862-y

Yarkoni, T., Poldrack, R. A., Nichols, T. E., Van Essen, D. C., & Wager, T. D. (2011). Large-scale automated synthesis of human functional neuroimaging data. Nature Methods, 8(8), 665– 670. 10.1038/nmeth.1635

Zhang, H., Zhao, R., Hu, X., Guan, S., Margulies, D. S., Meng, C., & Biswal, B. B. (2022). Cortical connectivity gradients and local timescales during cognitive states are modulated by cognitive loads. Brain Structure and Function, 227(8), 2701–2712. 10.1007/s00429-022-02564-0

Zhang, Y., Kim, J.-H., Brang, D., & Liu, Z. (2021). Naturalistic stimuli: A paradigm for multiscale functional characterization of the human brain. Current Opinion in Biomedical Engineering, 19, 100298. 10.1016/j.cobme.2021.100298

