## Supplemental Information for "Compression of functional gradients from rest to naturalistic processing: Moderation effect of age"

**Supplementary**

**Table S1. State differences in average gradient score of seven functional networks.**

|  |  | **Resting state** | **Movie watching** | ***t*-value** | ***p*-value** |
| --- | --- | --- | --- | --- | --- |
|  |  | **Mean±SD** | **Mean±SD** |  |  |
|  | **Can-CAM** |  |  |  |  |
|  | **Gradient 1** |  |  |  |  |
|  | VIS | -0.085 ± 0.002 | -0.071 ± 0.012 | -12.416 | <0.001 |
|  | SMN | 0.048 ± 0.012 | 0.045 ± 0.009 | 5.153 | 0.531 |
|  | DAN | 0.011 ± 0.009 | -0.003 ± 0.008 | 23.871 | <0.001 |
|  | VAN | 0.033 ± 0.009 | 0.024 ± 0.006 | 17.069 | <0.001 |
|  | LIM | 0.000 ± 0.009 | -0.001 ± 0.010 | 0.628 | <0.001 |
|  | FPN | 0.005 ± 0.006 | 0.006 ± 0.005 | -3.597 | <0.001 |
|  | DMN | 0.008 ± 0.006 | -0.003 ± 0.005 | -13.923 | <0.001 |
|  | **Gradient 2** |  |  |  |  |
|  | VIS | -0.039 ± 0.014 | -0.030 ± 0.008 | -11.914 | <0.001 |
|  | SMN | -0.044 ± 0.011 | -0.020 ± 0.009 | -37.644 | <0.001 |
|  | DAN | -0.024 ± 0.011 | -0.035 ± 0.008 | 17.454 | <0.001 |
|  | VAN | 0.015 ± 0.011 | 0.004 ± 0.007 | 17.168 | <0.001 |
|  | LIM | 0.027 ± 0.011 | 0.019 ± 0.010 | 10.168 | <0.001 |
|  | FPN | 0.031 ± 0.010 | 0.017 ± 0.006 | 24.538 | <0.001 |
|  | DMN | 0.049 ± 0.008 | 0.044 ± 0.005 | 10.891 | <0.001 |
|  | **Gradient 3** |  |  |  |  |
|  | VIS | -0.023 ± 0.008 | -0.009 ± 0.006 | -28.371 | <0.001 |
|  | SMN | -0.030 ± 0.009 | -0.027 ± 0.007 | -6.586 | <0.001 |
|  | DAN | 0.021 ± 0.008 | 0.018 ± 0.004 | 4.927 | <0.001 |
|  | VAN | -0.036 ± 0.009 | -0.014 ± 0.007 | -38.901 | <0.001 |
|  | LIM | 0.010 ± 0.011 | 0.006 ± 0.009 | 4.448 | <0.001 |
|  | FPN | 0.028 ± 0.008 | 0.025 ± 0.007 | 5.697 | <0.001 |
|  | DMN | 0.035 ± 0.009 | 0.012 ± 0.006 | 46.506 | <0.001 |
|  | **DyNAMiC** |  |  |  |  |
|  | **Gradient 1** |  |  |  |  |
| VIS | | -0.001 ± 0.011 | 0.001 ± 0.007 | -2.624 | 0.201 |
| SMN | | -0.057 ± 0.017 | -0.042 ± 0.016 | -9.832 | <0.001 |
| DAN | | -0.021 ± 0.010 | -0.025 ± 0.008 | 4.287 | <0.001 |
| VAN | | -0.024 ± 0.012 | -0.019 ± 0.009 | -4.431 | <0.001 |
| LIM | | 0.020 ± 0.011 | 0.010 ± 0.009 | 9.528 | <0.001 |
| FPN | | 0.031 ± 0.011 | 0.037 ± 0.011 | -4.538 | <0.001 |
| DMN | | 0.053 ± 0.012 | 0.037 ± 0.008 | 17.951 | <0.001 |
| **Gradient 2** | |  |  |  |  |
| VIS | | -0.061 ± 0.016 | -0.057 ± 0.010 | -2.888 | 0.093 |
| SMN | | 0.009 ± 0.008 | 0.011 ± 0.006 | -3.493 | 0.013 |
| DAN | | -0.010 ± 0.008 | -0.017 ± 0.007 | 9.058 | <0.001 |
| VAN | | -0.001 ± 0.009 | 0.004 ± 0.005 | -6.579 | <0.001 |
| LIM | | 0.013 ± 0.008 | 0.001 ± 0.007 | 4.707 | <0.001 |
| FPN | | 0.014 ± 0.007 | 0.018 ± 0.006 | -5.579 | <0.001 |
| DMN | | 0.032 ± 0.007 | 0.026 ± 0.005 | 9.813 | <0.001 |
| **Gradient 3** | |  |  |  |  |
| VIS | | 0.020 ± 0.009 | 0.017 ± 0.005 | 3.855 | 0.003 |
| SMN | | 0.019 ± 0.0011 | 0.016 ± 0.007 | 4.224 | <0.001 |
| DAN | | -0.026 ± 0.013 | 0.020± 0.007 | -4.941 | <0.001 |
| VAN | | -0.030 ± 0.011 | -0.020 ± 0.008 | -11.056 | <0.001 |
| LIM | | 0.006 ± 0.009 | 0.007 ± 0.007 | -1.730 | 0.624 |
| FPN | | -0.032 ± 0.010 | -0.028 ± 0.008 | -4.233 | <0.001 |
| DMN | | 0.016 ± 0.008 | 0.010 ± 0.006 | 8.154 | <0.001 |

Note: SD= standard deviation; VIS=visual network; SMN=somatomotor network; DAN=dorsal attention network; VAN=ventral attention network; LIM=limbic network; FPN=frontoparietal network; DMN= default mode network.

**Table S2. Association between functional integration/segregation, age and gradient metrics.**

|  | **PC** | | | | | | | | | | | **SS** |
| --- | --- | --- | --- | --- | --- | --- | --- | --- | --- | --- | --- | --- |
| Sparsity | 0.05 | 0.1 | 0.15 | 0.2 | 0.25 | | 0.3 | 0.35 | 0.4 | 0.45 | 0.5 | - |
| **Cam-can** | | | | | |  | | | | | | |
| ***Resting-state*** | | | | | |  | | | | | | |
| Age | 0.328** | 0.355** | 0.357** | 0.361** | 0.343** | | 0.334** | 0.326** | 0.319** | 0.313** | 0.308** | -0.260** |
| G1-range | -0.487** | -0.491** | -0.448** | -0.488** | -0.368** | | -0.337** | -0.311** | -0.291** | -0.274** | -0.257** | 0.338** |
| G2-range | -0.417** | -0.350** | -0.355** | -0.352** | -0.377** | | -0.380** | -0.383** | -0.381** | -0.378** | -0.373** | 0.331** |
| G3-range | -0.138* | -0.117* | -0.130* | -0.114* | -0.155* | | -0.158* | -0.160* | -0.160* | -0.157* | -0.153* | 0.346** |
| G1-variance | -0.507** | -0.519** | -0.476** | -0.515** | -0.393** | | -0.360** | -0.331** | -0.309** | -0.292** | -0.275** | 0.314** |
| G2-variance | -0.462** | -0.397** | -0.410** | -0.401** | -0.448** | | -0.456** | -0.461** | -0.462** | -0.459** | -0.454** | 0.292** |
| G3-variance | -0.060 | -0.059 | -0.068 | -0.053 | -0.083 | | -0.085 | -0.088 | -0.089 | -0.087 | -0.083 | 0.334** |
| ***Movie-watching*** | | | | | |  | | | | | | |
| Age | 0.319** | 0.315** | 0.280** | 0.307** | 0.195** | | 0.156* | 0.123* | 0.094 | 0.070 | 0.050 | -0.105* |
| G1-range | -0.252** | -0.283** | -0.301** | -0.298** | -0.286** | | -0.265** | -0.241** | -0.218** | -0.195** | -0.173** | 0.401** |
| G2-range | -0.201** | -0.162** | -0.127* | -0.092 | -0.059 | | -0.033 | -0.008 | 0.010 | 0.026 | 0.037 | 0.408** |
| G3-range | -0.185** | -0.124* | -0.071 | -0.025 | 0.011 | | 0.038 | 0.060 | 0.077 | 0.091 | 0.103 | 0.492** |
| G1-variance | -0.264** | -0.295** | -0.311** | -0.308** | -0.296 | | -0.274** | -0.249** | -0.226** | -0.204** | -0.181** | 0.398** |
| G2-variance | -0.374** | -0.348** | -0.314** | -0.279** | -0.246 | | -0.218** | -0.191** | -0.167** | -0.147* | -0.129* | 0.472** |
| G3-variance | -0.053 | 0.005 | 0.056 | 0.103* | 0.141* | | 0.167** | 0.189** | 0.205** | 0.216** | 0.226** | 0.410** |
| **DyNAMiC** |  |  |  |  |  | |  |  |  |  |  |  |
| ***Resting-state*** | | | | | | | | | | | | |
| Age | 0.226* | 0.252* | 0.244* | 0.233* | 0.221* | | 0.211* | 0.204* | 0.196* | 0.188* | 0.188* | -0.260** |
| G1-range | -0.683** | -0.660** | -0.658** | -0.664** | -0.670** | | -0.669** | -0.665** | -0.660** | -0.653** | -0.645** | 0.736** |
| G2-range | -0.437** | -0.438** | -0.438** | -0.422** | -0.401** | | -0.383** | -0.369** | -0.358** | -0.352** | -0.354** | 0.752** |
| G3-range | -0.470** | -0.429** | -0.408** | -0.393** | -0.387** | | -0.377** | -0.366** | -0.358** | -0.353** | -0.348** | 0.663** |
| G1-variance | -0.722** | -0.692** | -0.691** | -0.705** | -0.717** | | -0.723** | -0.727** | -0.728** | -0.726** | -0.720** | 0.653** |
| G2-variance | -0.427** | -0.416** | -0.419** | -0.410** | -0.396** | | -0.385** | -0.376** | -0.370** | -0.368** | -0.375** | 0.676** |
| G3-variance | -0.542** | -0.498** | -0.465** | -0.439** | -0.423** | | -0.405** | -0.389** | -0.377** | -0.368** | -0.362** | 0.608** |
| ***Movie-watching*** | | | | | | | | | | | | |
| Age | 0.506** | 0.500** | 0.473** | 0.455** | 0.442** | | 0.432** | 0.421** | 0.411** | 0.402** | 0.395** | -0.451** |
| G1-range | -0.591** | -0.615** | -0.634** | -0.643** | -0.648** | | -0.649** | -0.647** | -0.643** | -0.636** | -0.636** | 0.771** |
| G2-range | -0.436** | -0.472** | -0.491** | -0.500** | -0.499** | | -0.496** | -0.491** | -0.484** | -0.479** | -0.482** | 0.792** |
| G3-range | -0.427** | -0.335** | -0.285** | -0.247* | -0.221* | | -0.205** | -0.192* | -0.182* | -0.175* | -0.177* | 0.630** |
| G1-variance | -0.650** | -0.676** | -0.698** | -0.712** | -0.721** | | -0.725** | -0.726** | -0.725** | -0.720** | -0.720** | 0.724** |
| G2-variance | -0.513** | -0.541** | -0.555** | -0.557** | -0.555** | | -0.553** | -0.548** | -0.542** | -0.538** | -0.548** | 0.760** |
| G3-variance | -0.511** | -0.415** | -0.362** | -0.321** | -0.295** | | -0.279** | -0.266** | -0.256* | -0.250* | -0.256* | 0.590** |

Note: PC= participation coefficient; SS=system segregation. *: *p*<0.05; **: *p*<0.001.

**Table S3. Association between gradient compression and cognitive performance in Cam-CAN.**

|  | **Cognitive performance** | | | |
| --- | --- | --- | --- | --- |
|  | Cattell | Priming | Recognition | Recollection |
| ***With COV controlled*** | | | | |
| ΔG1-range | 0.077 | -0.020 | 0.138 | -0.009 |
| ΔG2-range | -0.076 | -0.106 | -0.220** | -0.159* |
| ΔG3-range | -0.077 | -0.121 | -0.090 | -0.146* |
| ΔG1-variance | 0.100 | -0.010 | 0.145* | -0.023 |
| ΔG2-variance | -0.025 | -0.013 | -0.145* | -0.155* |
| ΔG3-variance | -0.090 | -0.099 | -0.054 | -0.089 |
| ***With Age and COV controlled*** | | | | |
| ΔG1-range | 0.032 | -0.032 | 0.131 | -0.048 |
| ΔG2-range | 0.016 | -0.061 | -0.119 | -0.035 |
| ΔG3-range | 0.025 | -0.090 | -0.006 | -0.070 |
| ΔG1-variance | 0.040 | -0.028 | 0.120 | -0.030 |
| ΔG2-variance | 0.057 | 0.028 | -0.052 | -0.058 |
| ΔG3-variance | 0.001 | -0.083 | -0.006 | -0.046 |

Note: COV= covariance, including sex, education level, income, body mass index, total intracranial volume, and head motion. *: *p*<0.05; **: *p*<0.01.

**Table S4. Association between gradient compression and cognitive performance in DyNAMiC.**

|  | **Cognitive performance** | | |
| --- | --- | --- | --- |
|  | Episodic memory | Working memory | Perceptual speed |
| ***With COV controlled*** | | | |
| ΔG1-range | -0.121 | -0.054 | -0.126 |
| ΔG2-range | 0.008 | -0.033 | -0.115 |
| ΔG3-range | -0.028 | 0.033 | -0.108 |
| ΔG1-variance | -0.155 | -0.092 | -0.156 |
| ΔG2-variance | -0.060 | -0.028 | -0.060 |
| ΔG3-variance | -0.001 | 0.070 | -0.104 |
| ***With Age and COV controlled*** | | | |
| ΔG1-range | -0.070 | 0.025 | -0.058 |
| ΔG2-range | 0.058 | 0.025 | -0.075 |
| ΔG3-range | -0.003 | 0.082 | -0.095 |
| ΔG1-variance | -0.075 | 0.025 | -0.044 |
| ΔG2-variance | -0.081 | -0.018 | -0.059 |
| ΔG3-variance | 0.001 | -0.090 | -0.134 |

Note: COV= covariance, including sex, education level, body mass index, total intracranial volume, and head motion. *: *p*<0.05; **: *p*<0.001.

**Supplementary figures**

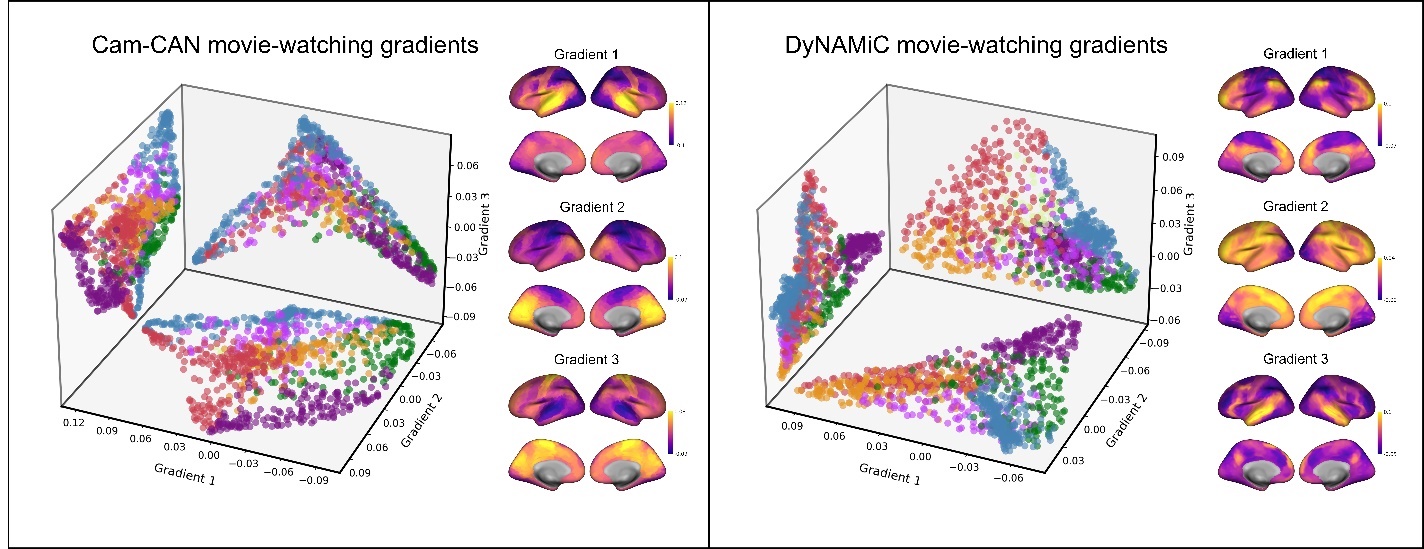

**Figure S1. The independent main movie-watching gradients.** In Cam-CAM, the first gradient is anchored by temporal areas, which are mainly responsible for auditory processing, and visual areas, forming a visual-auditory axis. For the second gradient, visual regions and DMN subregions such as the posterior cingulate cortex and precuneus are situated on one pole of the axis, while the sensory-motor areas and DAN areas gather at the other side. The third gradient, most transmodal and non-auditory regions have similar gradient scores, creating an auditory-nonauditory axis. In DyNAMiC, the first gradient shows a patten of S-A axis where DMN areas at one pole and unimodal areas (e.g., visual and sensorimotor areas) at the other pole. The second gradient, separates visual areas with other areas, making a visual-nonvisual axis. The third gradient anchored by auditory and language areas that located in temporal lobe, captured an auditory-nonauditory axis.

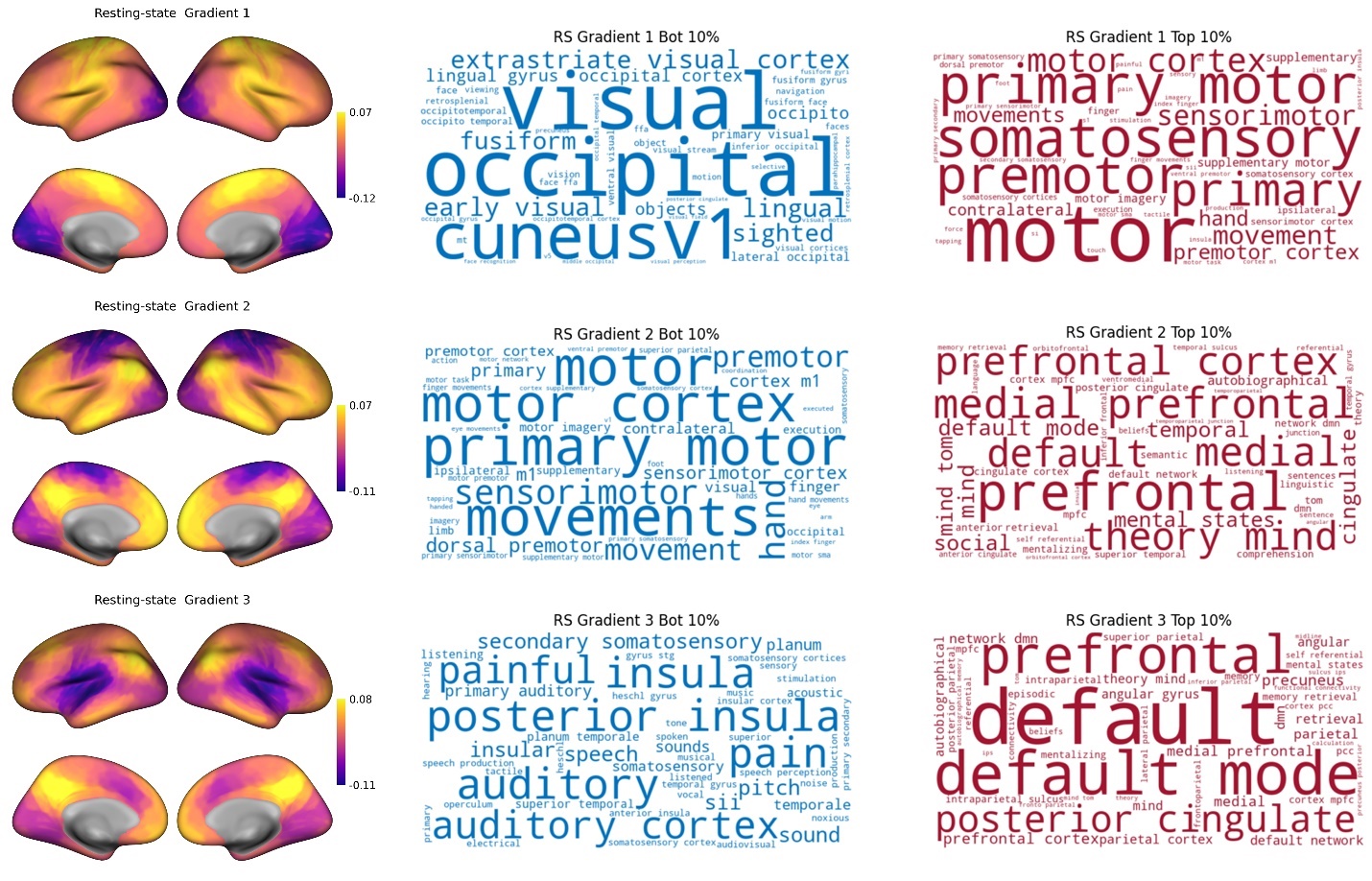

**Figure S2. Meta-analytic decoding of the first three main gradients in Cam-CAN.** Anatomical and functional terms generated from Neurosynth term-based decoding of the lowest and highest 10% of gradient scores per gradient. The font size of a given topic term represents the correlation between the gradient map and the meta-analytic map for that term, and a larger font size indicates a higher positive correlation.

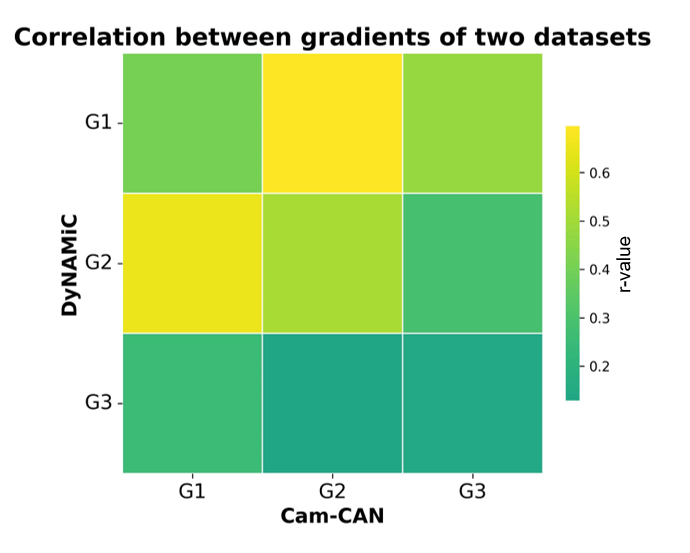

**Figure S3. The spatial correlations of the averaged resting-state gradient maps between two datasets.** The Cam-CAN G1 is highly correlated with DyNAMiC G2 (r = 0.653, p<0.001). The Cam-CAN G1 shows the pattern of the V-S axis, while DyNAMiC G2 forms a visual-nonvisual axis. Also, Cam-CAN G2 is strongly associated with DyNAMiC G1 (r = 0.696, p<0.001), as they follow the S-A axis.

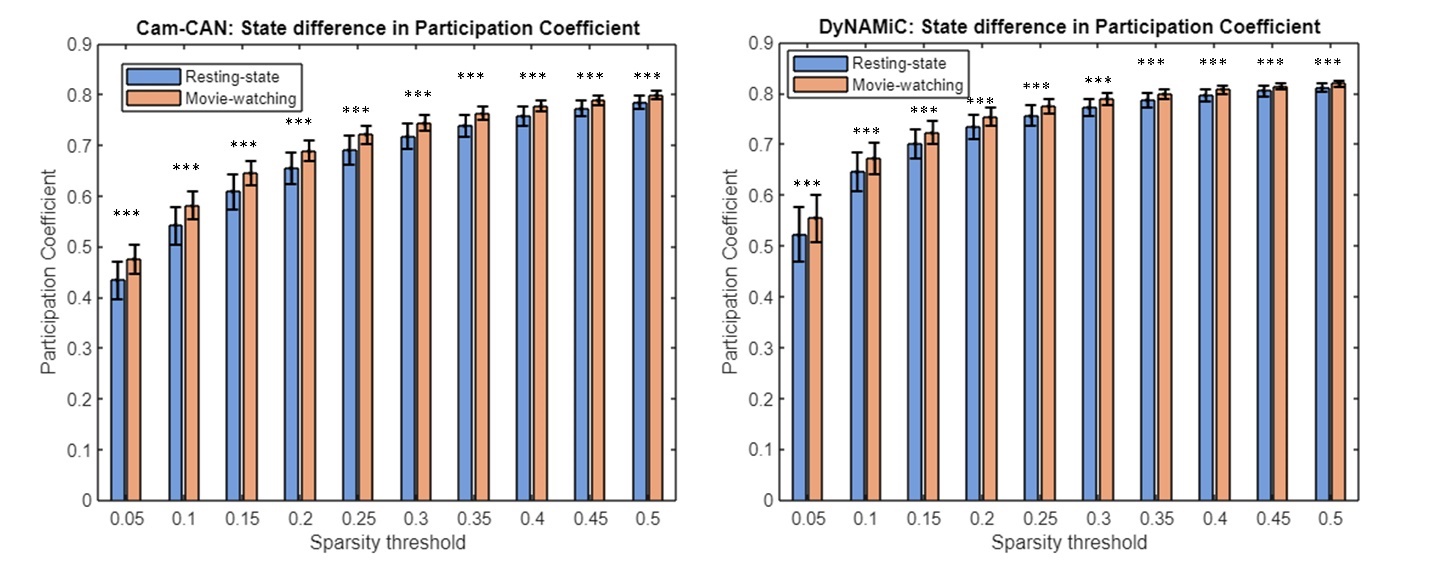

**Figure S4. The state difference in participation coefficients with different sparsity.** ***: *p*<0.001.

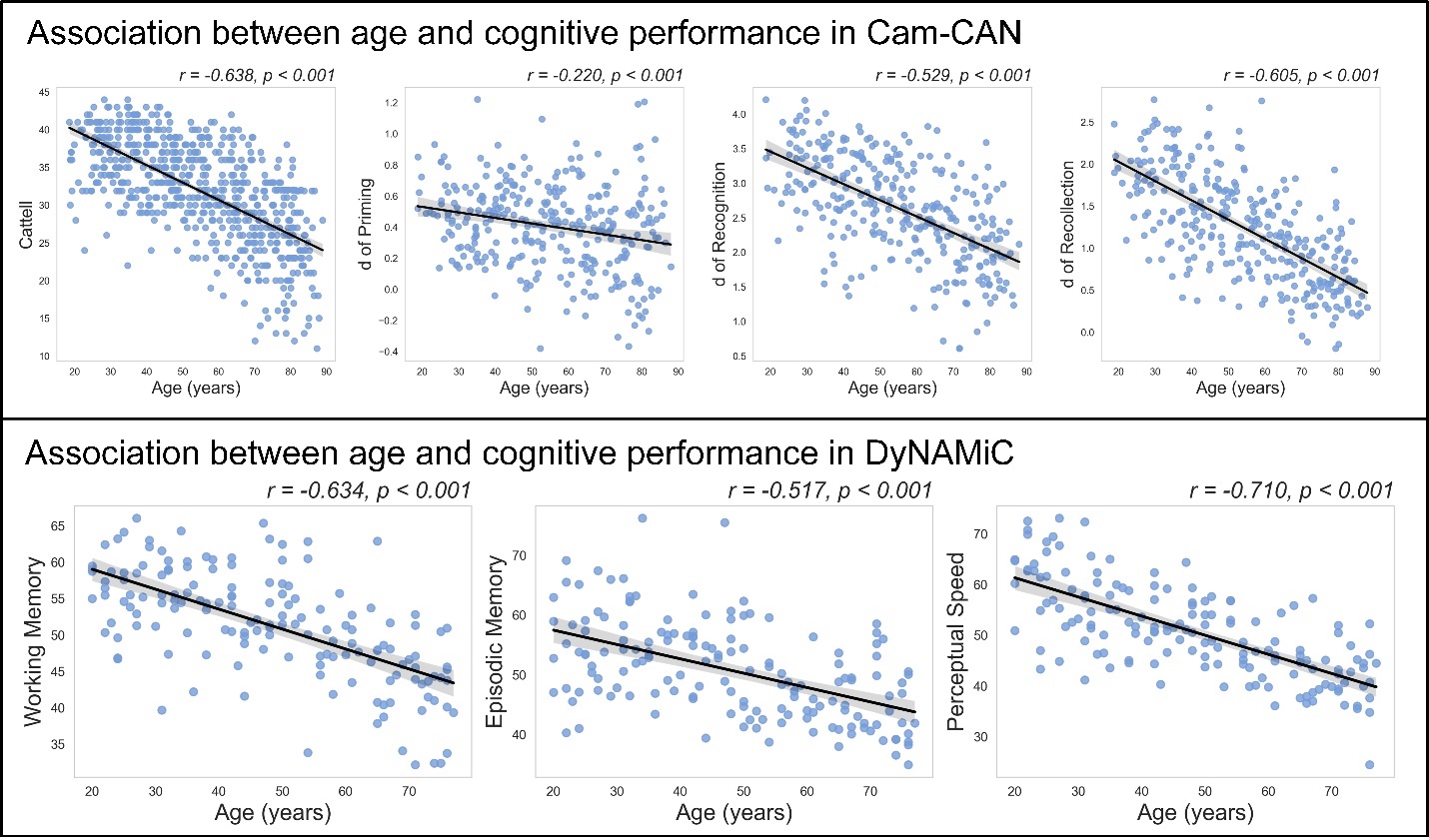

**Figure S5. Scatter plots of correlation between age and cognitive performance.**
